# XRN2-mediated regulation of tumor suppressor microRNAs is of critical pathophysiological significance in Humans

**DOI:** 10.1101/2021.08.20.457117

**Authors:** Rohit Nalawade, Tamoghna Chowdhury, Saibal Chatterjee

## Abstract

MicroRNAs (miRNAs) are critical regulators of diverse developmental and physiological processes, and they themselves get regulated both at the level of biogenesis and turnover. We demonstrate that the ribonuclease XRN2 can degrade the mature forms of certain let-7 family members in multiple human cancer cell lines, without affecting their precursors. XRN2 also affects the accumulation of several other tumor suppressor miRNAs known to play important roles in cancer metabolism. XRN2 depletion results in a reduction in the expression of many oncogenes and diminishes the proliferative and metastatic potential of cancer cells *in vitro*. These experimental cancer cells also show reduced capacity to form tumors in mice and regress over time. The clinical relevance of these observations is further verified in tumour transcriptomics data from public RNA-sequencing datasets, where *XRN2* mRNA expression is inversely correlated with the levels of a large number of miRNAs, including let-7 members, and high *XRN2* mRNA levels are associated with poor survival in hepatocellular carcinoma, lung adenocarcinoma, and glioblastoma. We demonstrate that the miRNA is released by an as-yet unidentified proteinaceous ‘miRNA release factor’ from the grasp of Argonaute before its degradation, which is more abundant in the nuclear fraction. Our analyses of the patient-derived transcriptomics data also show that XRN2, via its regulation of let-7, affects multiple pathways in a consistent manner across epithelial and glial cell lineages, and thus, is of critical pathophysiological significance.

## Introduction

miRNAs function as critical regulators of diverse developmental and physiological processes in animals^1^. Extensive studies on miRNA abundance and function have demonstrated a consistent link between dysregulation of miRNAs and various diseases, highlighting the importance of robust regulation of miRNA activity^2^. Many miRNAs exhibit tissue-and/or-stage-specific expression patterns^3^ and dramatic changes in the abundance of some miRNAs during developmental transitions of animals have also been reported^4^. Global kinetic studies of miRNA metabolism have revealed that regardless of high biogenesis, several miRNAs exhibit reasonably low steady-state levels^5,6^. It indicated that additional determinants, other than the factors of biosynthesis, play critical roles for the establishment of cellular miRNA homeostasis^5,6^.

A substantial amount of knowledge has been acquired on the different steps of miRNA biogenesis and their regulation, but much less is known about the miRNA turnover pathways and the constituent molecular machineries^7^. Active turnover of miRNAs in animals was first reported in *Caenorhabditis elegans* (*C. elegans*), where 5’-3’ exoribonuclease XRN-2 was demonstrated to mediate the degradation of several miRNAs^8^. Later, the paralogous protein, XRN-1, was also described as a ‘miRNase’ in worms^9,10^. In mammalian cells, several nucleases have been described to act on a given miRNA or a handful of miRNAs. XRN1 has been implicated in the turnover of miR-382 in HEK293 cells^11^, and polynucleotide phosphorylase degrades miR-221, miR-222, miR-106b in human melanoma cells^12^. Tudor-SN was demonstrated to target several functional mature miRNAs, including miR-31 and miR-29b, which control specific mRNAs encoding cell cycle regulatory proteins^13^. Interestingly, Argonaute1 (Ago1)-bound miRNAs in *Drosophila* are known to be tailed by terminal nucleotidyl transferase^14^ and trimmed by the 3’ – 5’ exoribonuclease Nibbler^15^ upon binding an exogenously introduced target that harbours extensively complementary binding-sites for the given miRNA^14,15^. Notably, in mice, an unusual extensively complementary endogenous target has been reported that lead to the 3’-end trimming-mediated destabilization of the cognate miRNA upon their interaction^16^. The terminal uridyl transferases, TUT4 and TUT7, are known to participate in tailing of miRNAs across species from *C. elegans* to humans^17^. A recent work using HEK293T cells has identified a machinery consisting of TUTs and DIS3L2, which could execute the decay of a subset of Argonaute (AGO)-bound mature miRNAs that have exposed 3’-ends^18^. Very recently, two simultaneous reports have substantially accentuated our understanding on the mechanisms of target-directed miRNA degradation (TDMD), where it was described that pairing of RISC/AGO-loaded miRNA to certain unusual highly complementary targets triggers ZSWIM8 ubiquitin ligase-mediated degradation of AGO, rendering the miRNA susceptible to decay by unknown nuclease(s)^19,20^. Perturbation of ZSWIM8 led to the accumulation of several miRNAs in different mammalian cells and some other systems (worms and flies). Intriguingly, half-lives of AGO proteins are much longer than majority of the miRNAs^5,6^, which suggested that ZSWIM8-mediated TDMD may not be the predominant miRNA turnover mechanism, rather, it is dedicated towards miRNAs that interact with unusual highly complementary targets. Thus, it is quite apparent that identification of yet unknown ‘miRNases’, followed by understanding of the pathways in which they are constituents, and their mechanistic details warrant further investigation.

Notably, although, knowledge about miRNA metabolism emanated from the studies employing worms have been found to be largely conserved, the role of the human ortholog of the worm ‘miRNase’-XRN-2 remains unclear. Apart from the critical roles played by XRN2 in multiple RNA-transaction pathways^21^, recent studies have also shown that it can act in an oncogenic capacity and worsen prognoses of specific cancers^22,23^. Here, we explore the capacity of human XRN2 as a ‘miRNase’ and elucidate the molecular mechanism(s) underlying the oncogenic capacity of XRN2 in relation to its ‘miRNase’ activity. In this regard, we focused on the fundamentally important let-7 family of miRNAs that are known to regulate important target mRNAs in multiple tissues, whose perturbation leads to different disease states^24^. We demonstrate that XRN2 specifically acts on the mature forms of most of the let-7 family members. Rapid and efficient depletion of XRN2 leads to the accumulation of AGO-bound let-7 members that in turn downregulate their cognate targets, including proto-oncogenes. Additionally, several other tumor suppressor miRNAs accumulate in XRN2-depleted cancer cells, whose oncogenic target mRNAs in turn show downregulation. Depletion of XRN2 in different cancer cells affect their cellular physiology by reversing critical cancer-parameters, including epithelial-to-mesenchymal transition. Pathophysiological significance of these observations got firmly established as the XRN2 depleted cancer cells not only show reduced capacity to form tumors in mice but also regress over time. Our *ex vivo* biochemical assays indicate that XRN2-mediated degradation of mature let-7 miRNAs happens upon ‘release’ of miRNAs from AGO by a nuclear enriched proteinaceous factor without affecting AGO integrity, and these two steps are kinetically linked, where miRNA function and turnover are largely spatially segregated in cytoplasm and nucleus, respectively. We also demonstrate that our cell line-based observations are of critical significance as they correlate with the clinical data of cancer patients from public transcriptomics datasets. Here, our analyses partially explain the multifarious roles of XRN2 in cancer through its likely participation in miRNA turnover, with the observation that XRN2 influences multiple pathways related to cellular proliferation, development, and signalling, especially related to extracellular matrix (ECM) development, and that these influences are mostly consistent across both epithelial and glial lineages. These analyses also partially explain previous observations of elevated *XRN2* mRNA expression being associated with worse survival in lung cancer patients and extend such observations into the more expansive Cancer Genome Atlas (TCGA) datasets for lung adenocarcinoma (LUAD), hepatocellular carcinoma (HPCC), and glioblastoma (GBM). Our analyses of these TCGA datasets also suggest that for many of the let-7 family members, *XRN2* demonstrates a strong inverse relationship comparable to some of the recently identified factors implicated in miRNA turnover, at an RNA level. Collectively, our study reveals that human let-7 miRNAs are regulated by a two-step turnover pathway, wherein, XRN2 plays the role of a ‘miRNase’ in various tissues.

## Results

### XRN2 affects only the mature forms of human let-7 family members

In human HEK293 cells, XRN1 has been implicated in the decay of miR-382, whereas depletion of the paralogous XRN2 had no significant effect^11^. The authors performed shRNA-mediated knockdown of *XRN1* and *XRN2*, but did not demonstrate the knockdown efficiencies at the protein level. Additionally, substantial knockdown of *XRN2* was not achieved in other related studies^11,25^, which could be insufficient to exert an effect on all the pathways XRN2 is involved in. Therefore, prior to investigating the role of XRN2 in miRNA degradation in human cells, we optimized a rapid and efficient shRNA-mediated specific knockdown of XRN2 using a lentivirus-based approach **(Supplementary Text 1, Fig. S1 a-d)**.

By means of turnover, XRN-2 in *C. elegans* modulates the mature forms of different members of the let-7 family, which play important roles in development. Dysregulation of their homologues also perturbs development and leads to many disease states in vertebrates^24^. Therefore, we began by examining the level of human let-7a miRNA, an identical homologue of *C. elegans let-7*, in control and XRN2 knockdown cells. In HEK293T cells, where the levels of endogenous let-7 miRNAs are very low^26^, XRN2 depletion did not affect the level of let-7a **(Fig. 1a)**. A possible reason for this could be that XRN2-mediated turnover might not be a predominant regulatory factor in maintaining its abundance in HEK293T cells. Contrastingly, ~3-fold accumulation in the level of mature let-7a was observed in XRN2-depleted A549 cells **(Fig. 1a)**.

**Figure 1.**
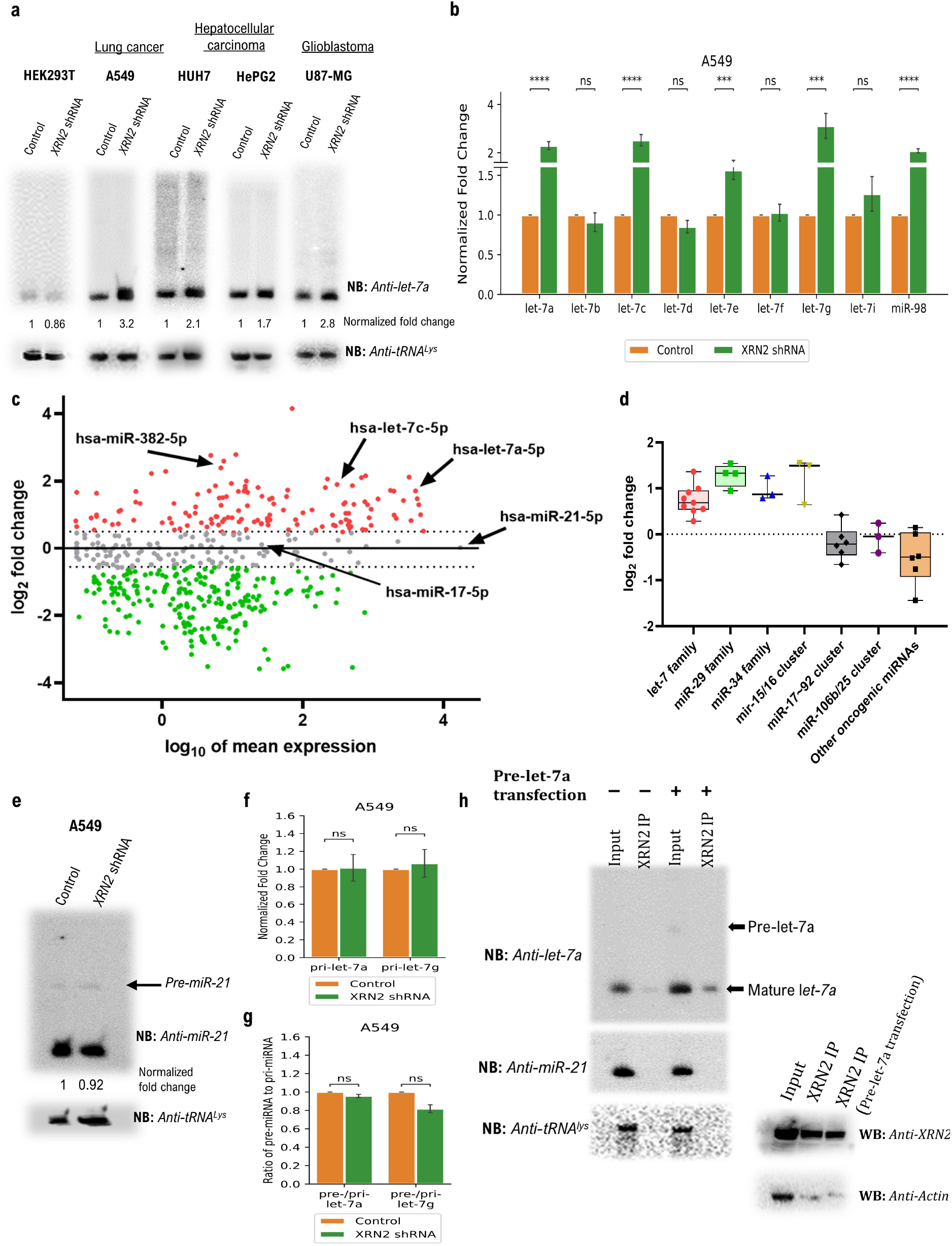
Accumulation of mature let-7a in different human cancer cell lines upon depletion of XRN2. **a.** XRN2 depletion leads to the accumulation of mature let-7a in A549, HUH7, HepG2, U87 cell lines, but not in HEK293T. **b.** TaqMan analyses reveal increased abundance of the mature forms of different let-7 family members upon depletion of XRN2 in A549 cell line. RNU48 RNA served as the normalization control. **c. Effect of XRN2 depletion on miRNA levels in A549 cells as measured by small RNA sequencing.** log2 fold change in miRNA expression (XRN2 shRNA over control) vs. miRNA expression in control. All the let-7 members that show upregulation in TaqMan assays also show 35% to ~2.5-folds upregulation in the small RNA-seq data of XRN2 depleted cells (compare **Fig. 1a-c, Fig. S2**). Oncogenic miRNAs (eg. miR-21-5p, miR-17-5p, indicated with arrows) do not show any appreciable change. Upregulated miRNAs are marked in red, while those downregulated marked in green. miRNAs in grey are the ones unchanged. **d. Box plot representation of small RNA-seq data from XRN2 depleted A549 cells reveal members of non-let-7 tumor suppressor family of miRNAs, as depicted, demonstrating comparable levels of accumulation to that of the let-7s.** Oncogenic miRNAs, as depicted, do not show accumulation. **e. Northern blot depicts unchanged abundance of the mature oncogenic miRNA-miR-21 in XRN2 depleted A549 cells.** Pre-miR-21 level also remains unaffected. **f.** RT–qPCR (n = 4; mean *±* SEM) analyses reveal no significant change at the level of primary RNAs for the indicated miRNAs in A549 cells. **g.** RT-qPCR (n = 4; mean *±* SEM) analyses depict no significant change in the relative levels of pre-miRNA transcripts for indicated miRNAs in A549 cells. **h. Left panel. Direct interaction between endogenous XRN2 and mature let-7a.** Co-immunoprecipitation reveals direct interaction between endogenous XRN2 and mature let-7a, but not the pre-miRNA. Non-substrate miRNA, miR-21, does not show any detectable interaction. **Right panel.** Anti-XRN2 western controls for the reactions depicted in the left panel.

In animals, let-7 miRNAs are known to exhibit different levels of abundance in different tissues, which could be determined by different regulatory forces^24^. Thus, turnover mediated regulation of specific miRNA levels might be more important in certain tissues, whereas regulated biogenesis might be more critical in others. Accordingly, we selected additional cell lines from different cell lineages, which are known to express adequately detectable levels of let-7 family members^27^. We incorporated glioblastoma (GBM) cell lines LN229 and U87-MG, hepatocellular carcinoma (HPCC) cell lines HUH7 and HEPG2, Breast cancer cell lines MDA-MB-231 and MCF-7, lung adenocarcinoma (LUAD) cell line A549 and Cervical cancer cell line HeLa in our study. XRN2-depleted lung (A549), hepatic (HUH7, HePG2), and neuronal (LN229, U87-MG) cells showed increase in the levels of their mature let-7a **(Fig. 1a, S2a)**. Conversely, let-7a miRNA did not accumulate upon XRN2 depletion in any of the breast or cervical cancer cell lines that we examined **(Fig. S2a)**.

Furthermore, we checked the effect of XRN2 depletion on other members of the let-7 family. TaqMan analyses revealed that XRN2 depletion affected many members, but not all. As demonstrated through Northern probing, TaqMan analyses also showed 2-3 folds accumulation of mature let-7a, while other members of let-7 family including let-7c, −7e, −7g, and miR-98 were also upregulated to different extents upon XRN2 depletion in A549, HUH7 and U87-MG cells **(Fig. 1b, S2c)**. However, let-7b, −7d, −7f, and −7i showed very little or no upregulation in the levels of their mature forms. Our northern and TaqMan assay-based observations were reiterated, when we analysed the small RNA sequencing data of control and XRN2 depleted A549 cells. Although, we did analyze only two sets of libraries that does not offer statistical validation, but still, all the let-7 members that showed accumulation in low-throughput experiments also showed 35% to ~2.5-fold upregulation in the high-throughput experiment (**Fig. 1c left panel**). We also noted that the members of other major tumor suppressor families of miRNAs like miR-29, miR-34, and miR-15/16 cluster were significantly upregulated in XRN2-depleted A549 cells, and comparable to the levels of upregulated levels of let-7s (**Fig. 1c right panel**). Importantly, We also checked the expression levels of miRNAs from several known oncogenic miRNA families like miR-21, miR-17~92, miR-106b/25, and miR-181 that have been shown to be important in the context of lung cancer. They remained unchanged upon XRN2 depletion (**Fig. 1c**), and could be re-confirmed through northern experiments as well (**Fig. 1d and data not shown**). Together, these data suggested that XRN2 can specifically affect certain tumor suppressor family members, including let-7, in a cell- or tissuespecific manner.

Additionally, we reckoned that being a ribonuclease, XRN2 could affect miRNA levels either by degradation of the mature forms or by regulating the processing of the corresponding precursor molecules during biogenesis. To answer this query, we determined the levels of the precursor RNAs of let-7a and let-7g, whose mature forms are most accumulated upon XRN2 knockdown. RT-qPCR analyses did not show any significant upregulation in the levels of the pri-let-7a and pri-let-7g **(Fig. 1c, S2d)**. The relative levels of pre-let-7a and pre-let-7g remained unchanged in the XRN2-depleted samples, in comparison to the control samples **(Fig. 1d, S2e)**. Accumulation of mature let-7a and let-7g without any appreciable change in the levels of any of their precursor forms indicated that XRN2 acts specifically on mature let-7 miRNAs leading to their degradation in A549, HUH7, and U87-MG cells. This inference was further strengthened as we could co-immunoprecipitate only the mature form of endogenous let-7a with endogenous XRN2, and not the pre-let-7a **(Fig. 1e)**. Even when cells were transfected with an exogenous pre-let-7a, more mature let-7 co-immunoprecipitated with endogenous XRN2, but still any association with pre-let-7a couldn’t be detected, although its abundant presence was well evident in the input sample. Furthermore, we also couldn’t detect any association of a non-substrate miRNA and endogenous XRN2 **(Fig. 1e**), which further confirmed that XRN2 not only acts on the mature forms, but also on specific substrate miRNAs.

### Accumulated miRNAs are functionally active: multiple validated targets of several tumor suppressor miRNAs get affected upon XRN2 depletion in multiple cancer cell lines

The tumor suppressor capability of let-7 miRNAs is attributed to the fact that they target several important genes including *RAS, HMGA2, MYC, CCND1*, and *CDC25A*^24^, which are potent oncogenes in humans. We anticipated that if the accumulated let-7 miRNAs in XRN2-depleted cells were not rendered non-functional, the levels of their target mRNAs should get affected, and that in turn would establish the regulatory role of XRN2 on the homeostatic capacity of let-7 on its cognate targets. Therefore, we compared the levels of the target mRNAs of let-7 family members (let-7s) in control and XRN2-depleted A549, HUH7, and U87-MG cells. Our mRNA sequencing (mRNA-seq) data from A549 samples revealed a considerable reduction in the mRNA levels of a large majority of the validated targets of let-7s (**Fig. 2a, c, Supplementary Table 1**). Moreover, cumulative distribution function (CDF) of mRNA changes for predicted and validated targets of let-7s derived from the mRNA-seq results revealed that the median values of let-7-targets were less than the median of the non-targets, as it would be expected of let-7-targets being downregulated upon XRN2 depletion. The non-target empirical CDF more closely followed a random normal distribution than the target empirical CDF (**Fig. 2b**), which clearly suggested that the targets were more likely regulated directly rather than being affected by chance factors. We also noted promising results from the CDF analyses performed with mRNA-seq data obtained from HUH7 and U87 samples **(Fig. S3a, b,** and **data not shown**). Further, for the sake of brevity, we have highlighted only one and half dozen validated pathophysiologically important let-7-targets that were significantly downregulated in XRN2-depleted A549 cells (**Fig. 2c**). These regulatory effects observed on let-7 cognate targets could also be confirmed by low-throughput RT-qPCR methods, and they by and large supported our inferences made from the mRNA-seq data (**Fig. 2d, S3c, d**). Additionally, mRNA-seq data also revealed downregulation of validated targets of the non-let-7 tumor suppressor miRNAs that were accumulated, and the effects they exerted on their targets were comparable to that of the let-7s (**Fig. 2e**). Notably, the validated tumor suppressor mRNA-targets of the known oncogenic miRNAs, which remained unchanged, were not downregulated in the experimental datasets (**Fig. 2e**), and thus there was no potential nullification of the effects of downregulation of oncogenic targets mediated by the accumulated tumor suppressor miRNAs.

**Figure 2.**
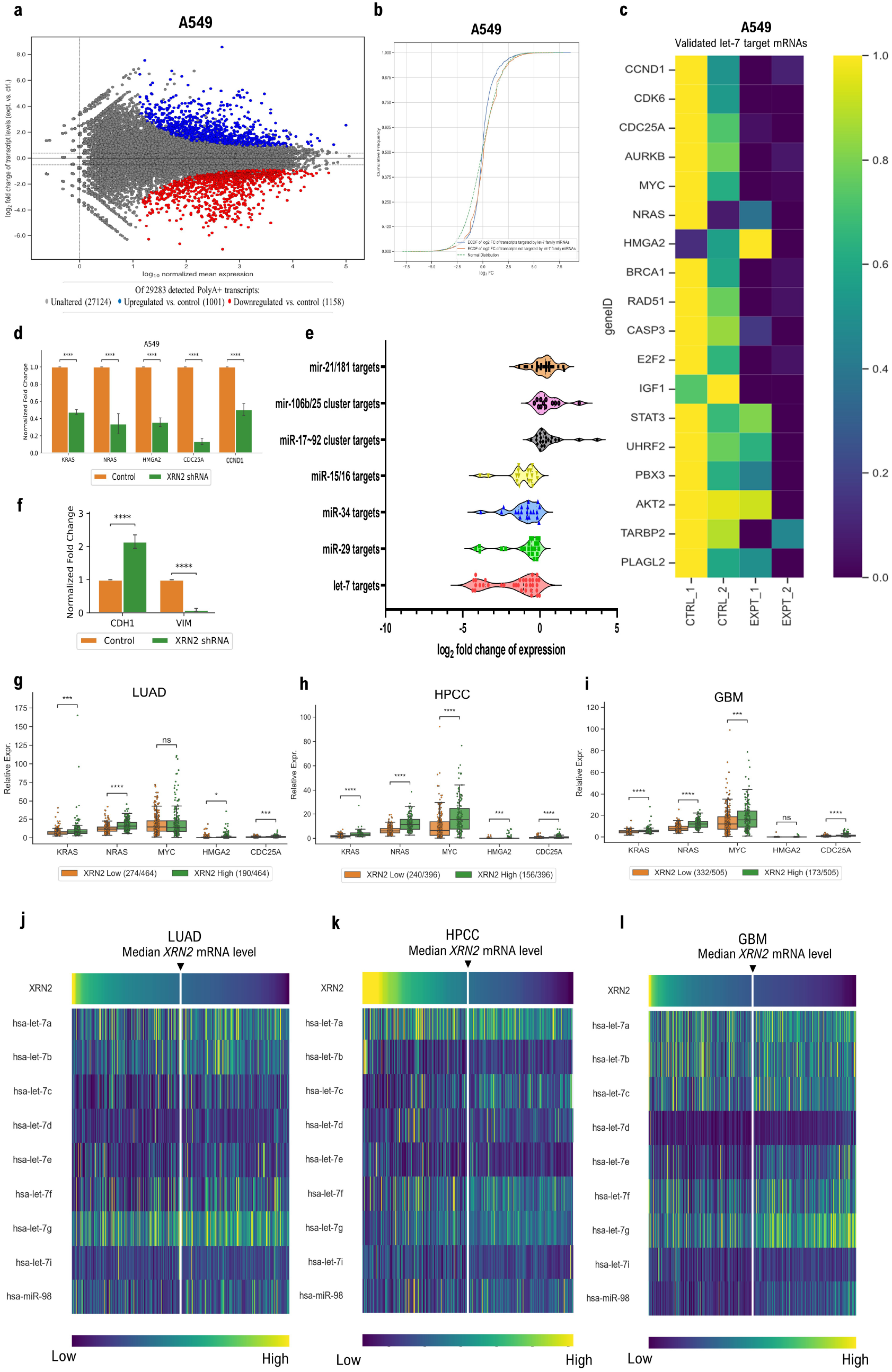
XRN2 depletion results in the downregulation of key let-7 and other tumor suppressor miRNA targets in cancer cell line and increased *XRN2* mRNA levels positively affect expression of various oncogenes and negatively influence the expression of let-7 family members in patients. **a.** MA/ Bland-Altman plot of expression of detected polyA+ transcripts for annotated genes. In red are genes whose transcripts are downregulated at least *0.7 ×* with respect to the control samples (1158/29283, 3.9%), and in blue are genes whose transcripts are upregulated at least 1.3 *×* with respect to the control samples (1001/29283, 3.4%). **b.** Cumulative distribution function (CDF) of mRNA changes for predicted let-7 target (depicted in blue) and non-target (depicted in red) genes indicates downregulation of let-7 targets. **c.** Relative mRNA expression levels (arbitrary units) of the indicated let-7 target genes known to act as oncogenes in the control and XRN2 knockdown samples from A549 cells. **d.** RT–qPCR (n = 4; mean *±* SEM) analyses reveal reduction in let-7 target mRNAs, as indicated, upon XRN2 depletion in A549 cells. Fold changes were normalized to *ATP5G* mRNA. P-values are from paired Student’s t-test. **e. Violin plot representation of mRNA-seq data from XRN2-depleted A549 cells reveal targets of non-let-7 tumor suppressor family of miRNAs, as depicted, showing down regulation comparable to that observed for the targets of let-7s.** Targets of oncogenic miRNAs (tumor suppressor mRNAs), as depicted, do not show downregulation. **f.** RT–qPCR (n = 4; mean *±* SEM) analyses reveal changes in the mRNA levels of EMT genes, as indicated, upon XRN2 depletion in A549 cells. Fold changes were normalized to *ATP5G* mRNA. P-values are from paired Student’s t-test. **g-i.** Relative mRNA expression levels (arbitrary units) of the indicated let-7 target genes in the high and low *XRN2* mRNA expression groups from the TCGA-LUAD (**left panel**), TCGA-HPCC (**middle panel**), TCGA-GBM (**right panel**) RNA-Seq datasets of clinical LUAD, HPCC, GBM tumour samples, respectively. P-values are from Student’s t-test with Bonferroni correction. **j-l.** Expression patterns of let-7 family members in relation to *XRN2* mRNA levels in the TCGA-LUAD (**left panel**), TCGA-HPCC (**middle panel**), TCGA-GBM (**right panel**) RNA-Seq datasets. let-7a, −7c, −7f and −7g are visibly inversely correlated with *XRN2*. Colour Gradient: Yellow – highest expression, Blue – lowest expression (relative to the minimum and maximum expression of the respective miRNA or mRNA). The split line through the heatmap denotes the median expression value of *XRN2* mRNA.

To further enrich our observations, we also checked the levels of Epithelial-to-Mesenchymal transition (EMT) genes in these samples as certain let-7 family members have already been shown to function as vital regulators of the EMT. As expected, depletion of XRN2 in A549 cells, drastically reduced the *VIM* mRNA expression, while the expression of *CDH1* mRNA showed a significant upregulation in both RT-qPCR analysis and mRNA-seq data (**Fig. 2f, S3e**). Of note, a reporter gene hosting let-7a target sites in its 3’-UTR showed downregulation in XRN2 depleted cells. This regulatory effect was disrupted when those let-7a target sites were deleted, which essentially confirmed that the regulatory effect of accumulated let-7a got exerted through a direct sequence-based interaction between the miRNA and the target site (**Fig. S3f**). Thus, in light of all the above observations, it could be suggested that to a reasonably large extent the effect of depletion of XRN2 on the transcriptome was indeed due to its role as a ‘miRNase’, which further advocated that the accumulated let-7 family members were functionally active.

To augment our observations derived from cell culture-based studies, we investigated if the mRNA levels of the validated let-7 targets *KRAS, NRAS, MYC, HMGA2* and *CDC25A* were also accordingly affected in patients displaying high expression of *XRN2* mRNA in a public transcriptomics TCGA dataset for HPCC, LUAD and GBM. Each of these mRNAs was elevated in patients having higher than median *XRN2* expression as compared to patients with lower than median expression of *XRN2* in all three datasets (**Figure 2g-i**). Notably, we employed median values as they get less affected by positive/ negative skewing of heterogeneous data, as well as by outliers than the mean values. The above observations indeed corroborated with our results obtained from the aforementioned XRN2-depleted cell lines. Having observed this dysregulation of the oncogenic let-7 targets, we then investigated the relationship of let-7 family members with *XRN2* mRNA levels in the same TCGA datasets. Based on the heatmaps presented in **Fig. 2j-l**, which indicated that let-7a, let-7c, let-7f, let-7g, and miR-98 depict an inverse relationship with *XRN2* mRNA expression (**S4a-c, Supplementary text 2)**, we chose to perform all subsequent analyses on a randomly-selected subset of the TCGA-HPCC, TCGA-GBM, and TCGA-LUAD datasets broadly possessing the greatest combined mean fold change for let-7c and let-7g for low over high *XRN2* mRNA expression, as well as the strongest combined inverse correlation of let-7c and let-7g with *XRN2* mRNA expression (see Methods for detailed selection procedure), hereafter referred to as a ‘representative subset’. We expected that this approach would allow us to select a set of samples, where XRN2-mediated regulation of let-7 miRNAs was optimally operational for further comparative studies. These data indicated that the specificity of XRN2-mediated regulatory activities vary in different clinical tissue samples, similar to the way it varied in the above-described defined cancer cell lines (compare **Fig. 2g-i** with **Fig. 2d, S3c, d**).

Outcomes of the aforementioned analyses of the patient samples intrigued us to investigate whether the effect of XRN2 extends beyond the family of let-7 miRNAs. Accordingly, we compared the levels of other miRNAs with *XRN2* mRNA in the abovementioned patient-derived datasets. Around 10% of all the cellular miRNAs, including some of the highly conserved and functionally important ones, showed a significant inverse relationship with *XRN2* mRNA levels in the ‘representative subsets’ from all three datasets (‘representative subsets’ for TCGA-GBM and TCGA-LUAD were obtained following the same criteria and process described for TCGA-HPCC) (**Fig. S4d-f**). To validate and complement our *in silico* observations, we selected one ‘potential non-let-7 XRN2-substrate’ miRNA from each dataset, and checked its level in a cell line of the respective lineage upon XRN2 depletion. We assessed the levels of miR-122 in HUH7 cells (HPCC), miR-124 in U87 cells (GBM), and miR-29a in A549 cells (LUAD)^2,31^. Indeed, upon XRN2 depletion, we observed significant accumulation of all the three tissue specific miRNAs in the respective cell lines (**Fig. S4d-f**). These observations and results indicated that XRN2 has a significant impact on the total miRNA pools in different cell lineages and therefore, XRN2-mediated miRNA turnover might constitute an important pathway in cellular miRNA metabolism.

### Effects of XRN2 depletion on cancer cells

let-7 miRNAs are known to repress proliferation and facilitate differentiation by affecting multiple critical pathways in human cells^24^. Previous studies in human LUAD and ovarian cancer cell lines have reported that overexpression of certain let-7 family members results in anti-proliferative effects and reduced survival of cancer cells^24^. Our observations on the levels of mature let-7 miRNAs upon XRN2 knockdown, with a concomitant effect on the levels of crucial target-oncogenes, prompted us to check if XRN2 depletion can lead to the reversal of tumor phenotypes in these cancer cell lines. Notably, cell migration is a crucial process in cancer progression that allows cancer cells to metastasize to different tissues through the circulatory system. To test whether XRN2 depletion can affect the migration of these cancer cells, we performed a wound healing assay, which allowed us to study directional migration of cancer cells *in vitro*. We observed that the migration of A549, HUH7 and U87-MG cells undergoing XRN2 knockdown was significantly reduced as compared to the control cells. Interestingly, introduction of a specific let-7a antagomir, and not a non-specific sequence, to those cells undergoing XRN2 knockdown reversed the inhibitory effect on migration, which firmly suggested that the effects were indeed mediated by the accumulated let-7s through sequence specific interactions **(Fig 3a, b, S5a, b, S5e, f)**. We further performed clonogenic assays, which examines the ability of a single cancer cell to undergo multiple divisions and grow into a colony. This assay performed with all the three cell lines revealed diminished colony formation ability in XRN2-depleted cells as compared to the control cells **(Fig 3c, d, S5c, d, S5g, h)**.

**Figure 3.**
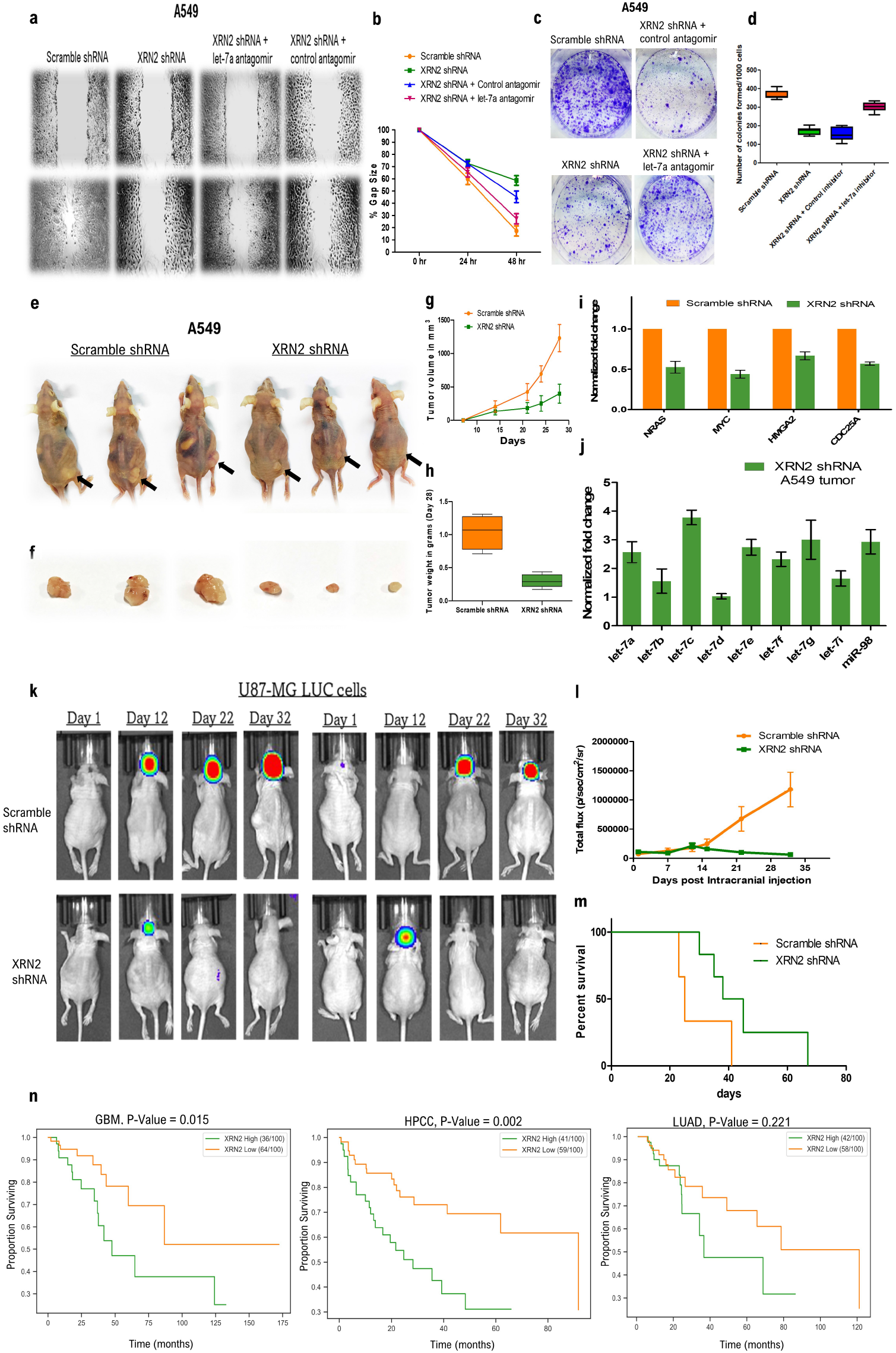
XRN2 depletion affects physiology of cancer cells *in vitro* as well as in xenograft experiments, and high *XRN2* mRNA expression worsens survival of cancer patients. **a, b.** Wound-healing assay performed with A549 cells (n=4) transfected with indicated shRNAs depict reduced cell migration in cells transfected with XRN2 shRNA compared to scramble shRNA. Treatment with let-7a antagomir efficiently reverses decreased cell migration of XRN2-depleted A549 cells. Graphical representation of the wound-healing assay reveals significant differences in the gap-size between the indicated samples. **c, d.** Clonogenic assay performed with A549 cells employing scramble shRNA and XRN2 shRNA (n=4) depicts decreased colony formation ability of XRN2-depleted cells. Corresponding box plot demonstrates the decrease in the number of colonies formed by XRN2-depleted cancer cells and reversal of the phenotype upon subjecting let-7a antagomir treatment to the XRN2-depleted cells. **e, f.** Representative images of mice injected subcutaneously with either scramble shRNA or XRN2 shRNA transfected A549 cells after 28 days of injection. **(g, h)** Graph representing the tumor volume (mm^3^) and tumor weight in grams (gm) for scramble shRNA and XRN2 shRNA transfected A549 tumors after 28 days of injection. **i.** RT-qPCR analyses reveal downregulation of expression of let-7 target mRNAs in XRN2-depleted A549 tumors compared to control tumors. RT-qPCR (n = 5; mean ± SEM) results were normalized against ATP5G mRNA. **j.** TaqMan analyses reveal increased abundance of the mature forms of different let-7 family members in XRN2-depleted A549 tumors compared to control tumors. **k.** *In vivo* whole-body bioluminescent imaging of mice injected intracranially with either scramble shRNA transfected U87-MG LUC cells or XRN2 shRNA transfected U87-MG LUC cells depict drastic differences in the growth patterns of tumors over time. **l.** Quantification of the total radiance of bioluminescent signals depicted in k. The two groups of animals are indicated with different colors, as indicated. ANOVA was used to calculate the significance of differences between the groups. **m.** Kaplan-Meier graphs demonstrate the difference in the survival of mice bearing intracranially injected U87-MG LUC cells transfected with either scramble shRNA or XRN2 shRNA. **n.** Kaplan-Meier survival analyses of TCGA datasets indicate association of high XRN2 mRNA levels with worsened survival outcomes in glioblastoma (TCGA-GBM), hepatocellular carcinoma (TCGA-HPCC), and lung adenocarcinoma (TCGA-LUAD). p-values are from Chi-Square test with d.f. = 1 (2 categories).

To further assess and understand the effect of XRN2 depletion on the tumorigenic properties of these cancer cells from an *in vivo* perspective, we resorted to xenograft experiments. We planted those control and knockdown cells subcutaneously in nude mice and followed the fate of those cells **(Fig 3e, S6a)**. The control cells exhibited uninterrupted growth and proliferation and resulted in the formation of large tumors. Contrastingly, the knockdown cells resulted in the formation of much smaller cell masses and showed signs of regression during certain phases of the time course **(Fig 3f-h, S6b-d)**. Notably, the tumors formed from the knockdown cells, compared to the control cells, showed higher expression of tumor suppressor miRNAs (eg. let-7s) and lower expression of their cognate oncogenic target mRNAs (**Fig. 3i, j, S6e, f)**, which in turn indicated that those XRN2/let-7s-regulated target mRNAs are indeed important for the growth of the tumors. Intrigued by these results, where the cancer cells were placed in ectopic locations, we implanted xenografts with U87-MG LUC cells by placing them in *bona fide* location, the intracranial chamber of nude mice. Mice implanted with control cells formed large tumors that continued to grow till the last recorded time point. Conversely, mice implanted with XRN2 knockdown cells formed tumorous cell masses during the earlier time points but got regressed over time **(Fig. 3k, 3l)** and no tumorous cell mass could be detected in histological tissue sections **(data not shown)**, which eventually resulted in their significantly longer survival than the mice implanted with control cells **(Fig. 3m)**. Overall, these results demonstrated that the changes in miRNA levels upon XRN2 depletion were indeed physiologically significant as they affected the tumorigenicity of the cancer cells.

As our above-mentioned results demonstrated XRN2-mediated regulation of let-7 miRNAs in cancer cells from LUAD, HPCC, GBM (**Figs. 1a-e, S2c**) as well as negative correlation between *XRN2* mRNA and let-7 members (**Fig. 2i-k**), we further performed an independent Kaplan-Meier survival analysis on the ‘representative subsets’ of the heterogeneous public datasets for TCGA-HPCC, TCGA-GBM, and TCGA-LUAD to explore any possible link of patient lifespan with *XRN2* expression in these cancers. Indeed, these analyses further verified that elevated levels of *XRN2* mRNA were associated with worsened survival outcomes in the patients who contributed to these datasets **(Fig. 3n)**.

### Turnover of let-7 miRNAs is kinetically linked to the release of miRNAs bound to AGO2

miRNA biogenesis is completed with the generation of an AGO-loaded mature miRNA, as a constituent of a macromolecular complex called miRNA-induced silencing complex (miRISC)^1^. We wanted to check how XRN2 depletion affects the mature miRNAs bound to the AGO proteins. Accordingly, we immunoprecipitated the endogenous AGO2 from either control or XRN2-depleted A549 cells to check the levels of let-7a miRNA associated with immunoprecipitated AGO2. A significantly increased signal was recorded from AGO2 immunoprecipitated from the XRN2-depleted cells **(Fig. 4a)**. Similar observations were made upon immunoprecipitation of AGO2 from HUH7 and U87 cells **(Fig. S7a)**. Furthermore, we checked the levels of other let-7 sisters associated with immunoprecipitated AGO2 from A549 cells. Here, we observed that amongst all the let-7 miRNAs, let-7a, −7c, −7e, −7g, and miR-98 showed the highest accumulation in immunoprecipitated AGO2 from XRN2-depleted cells compared to control (**Fig. 4b**), and that further indicated to a dislocation in the cascade of events leading to turnover of these miRNAs. We were intrigued by this observation to check whether those miRNAs undergo a ‘release step’ from AGO before degradation, and how XRN2 may affect this process. Accordingly, we performed an *ex vivo ‘*miRNA release assay’^8^, employing immunoprecipitated endogenous AGO2 from A549 cells **(Fig. 4c)**. Immunoprecipitated bead-bound AGO2, upon incubation with control lysate and a further recovery, showed a significant loss of let-7a signal, when compared to the signal obtained from the bead-bound AGO2 incubated with buffer **(Fig. 4d)**. No miRNA could be detected in the RNA recovered from the supernatant fraction of the release assay, where bead-bound immunoprecipitated AGO2 was treated with control lysate **(Fig. 4d)**. However, the assay that employed XRN2-depleted cell lysate for incubation with bead-bound immunoprecipitated AGO2, more let-7a could be detected in the recovered bead-bound immunoprecipitated AGO2, post treatment **(Fig. 4d)**. A substantial amount of let-7a could also be detected from the RNA extracted from the supernatant fraction of the same reaction **(Fig. 4d)**, which clearly demonstrated that those miRNAs are not associated with AGO2, but still not degraded. This data suggested that a diminished level of XRN2 in knockdown samples might be good enough to perform the ‘release activity’ but unable to degrade the released let-7a miRNA. Alternatively, XRN2 is kinetically linked to the process of let-7a release from AGO2, performed by a dedicated ‘miRNA release factor’ that is not present in molar excess in comparison to XRN2. This latter notion was strengthened as we observed that the ‘XRN2-non-substrate’ miRNA, miR-21, got released equally well by either of the control or XRN2 knockdown lysate **(Fig. 4d)**. Importantly, the released miR-21, unlike let-7a, didn’t get protected in the supernatant fraction of the reaction involving the XRN2 knockdown lysate, which not only suggested that some other factor(s) led to the degradation of miR-21, but also reconfirmed the specificity of XRN2. However, these results didn’t resolve the question whether same or different ‘release factors’ took part in the releasing of XRN2-substrate and non-substrate miRNAs. Additionally, since, no XRN2 protein could be detected with immunoprecipitated AGO2 **(Fig. S7b)**, it could be suggested that either the AGO2-XRN2 interaction is highly transient or there is indeed a yet unidentified ‘miRNA release factor’ that facilitates miRNA release from AGO2 for its degradation. Notably, subjecting the above-mentioned control cell lysate with protein denaturing conditions resulted in a complete loss of its ‘miRNA releasing activity’, which suggested a proteinaceous nature of the ‘miRNA release factor’ **(Fig. S7c)**.

**Figure 4.**
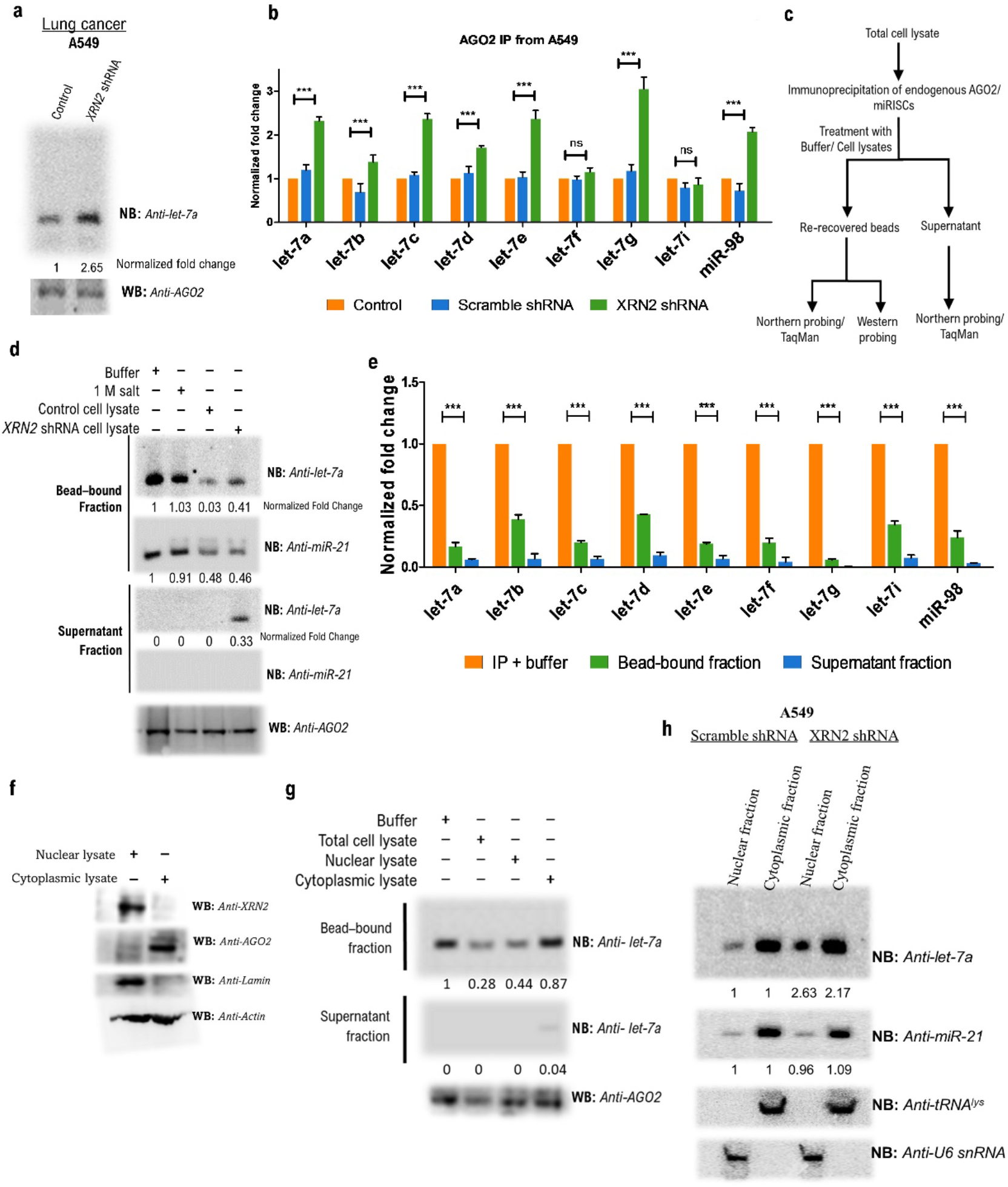
Increased signal of let-7 miRNAs from AGO2 immunoprecipitated from XRN2-depleted cells. **a.** Immunoprecipitation using anti-EIF2C2 (anti-AGO2) antibody from total cell lysate of the indicated samples reveal increased let-7a signal from AGO2 of XRN2-depleted cells. One half of each immunoprecipitate was subjected to northern probing or TaqMan analysis, the other half to anti-AGO2 western probing and served as loading control. **b.** TaqMan analyses (n = 3; mean ± SEM) reveal increased accumulation of several let-7 sisters in immunoprecipitated AGO2 from XRN2-depleted samples. **c.** Schematic representation of the miRNA release assay. Bead-bound immunoprecipitated AGO2/ miRISC subjected to different treatments as indicated and recovered back as bead and supernatant fractions. One half of each re-recovered bead fraction is subjected to northern probing or TaqMan analysis, the other half to anti-AGO2 western probing. Northern probing or TaqMan analysis is performed for equivalent amount of RNA extracted from the supernatant fraction. Loss of signal from the re-recovered intact AGO2/ miRISC subjected to a specific treatment, compared to that of the buffer treatment, indicates release of AGO2/ miRISC-bound let-7a miRNA. **d.** miRNA release assay with immunoprecipitated AGO2 was performed with the indicated samples. Release of let-7a is diminished in the assay employing XRN2-depleted total lysate compared to the one with control lysate. **e.** TaqMan analyses (n = 4; mean ± SEM) of the RNA isolated from re-recovered bead-bound fraction reveal differential release of different let-7 sisters from immunoprecipitated AGO2 subjected to control cell lysate. The supernatant fraction depicts less than 5% of the miRNA signals obtained from the bead-bound fraction of the control reaction. **f.** Nucleo-cytoplasmic distribution of XRN2 and AGO2 in A549 cells. Lamin serves as nuclear marker. **g.** Nuclear fraction is more enriched with the miRNA release activity compared to that of the cytoplasmic fraction. **h.** XRN2 knockdown leads to a greater fold-change of nuclear localized let-7a compared to that localized in cytoplasm. Relative nucleo-cytoplasmic distribution of the non-substrate miRNA (miR-21) remains unchanged upon XRN2 knockdown.

As treatment with total cell lysate resulted in an efficient release of let-7a from immunoprecipitated AGO2, we checked whether other let-7 sisters got similarly released from the AGO2 protein. We observed that let-7a and let-7g underwent release to the extent of 85-90%, while the other sisters showed 60-80% release from AGO2 **(Fig. 4e)**. We also measured let-7a and other sisters from the supernatant fraction, and they could only be minimally detected, accounting for less than 5% of the signal obtained from buffer-treated immunoprecipitated bead-bound AGO2 **(Fig. 4e).**

### Nuclear fraction is enriched with miRNA releasing and turnover activities

The above results on release of miRNAs from AGO2/miRISC before their degradation raised an obvious question about the site(s) of miRNA release and degradation. This is intriguing as XRN2 is known to be predominantly nuclear^21^, whereas cytoplasm is the work bench for miRISC function. Notably, recent reports from multiple systems have shown considerable to modest nuclear localization of Argonaute proteins^51^, where they perform certain roles^51^. Accordingly, upon nucleo-cytoplasmic fractionation of A549 cells, we observed a predominantly nuclear localization for XRN2. 10-15% of AGO2 and a similar proportion of let-7a could be detected in the nuclear fraction (**Fig. 4f, 4h**). We checked the miRNA releasing activity of these two fractions and noted that the nuclear fraction has a much stronger releasing activity than the cytoplasmic one (**Fig. 4g**). This observation suggested that the nuclear localized AGO2/miRISC are more likely to encounter the ‘release factor’ leading to the turnover of their resident miRNAs. Thus, the nuclear compartment potentially acts as a prime site of turnover that is spatially segregated from the site of miRNA function (cytoplasm). However, this possibility by no means disfavours AGO2/miRISC to perform a specific function in the nucleus. On the other hand, the modest cytoplasmic miRNA releasing activity could be a part of a surveillance mechanism, ensuring the release of a miRNA from AGO2/miRISC that is not engaged in an interaction with its cognate target mRNA, and thus allowing the ‘freed’ AGO2 to be reloaded with a new miRNA.

Interestingly, we observed a greater fold-change of let-7a accumulation in the nuclear fraction compared to the cytoplasmic one upon knockdown of XRN2. But the quantity of accumulation in the cytoplasmic fraction was much higher compared to the nuclear fraction. These observations suggested that the miRNA release factor is not likely to be in molar excess in the cytoplasm and might travel to nucleus for the delivery of its ‘cargo’ to the ‘miRNase’, where depletion of the ‘miRNase’ perturbs the kinetics of its sequential interactions and roles. This notion corroborates with our aforementioned results, where we presented increase in AGO2 associated let-7a and its activity upon depletion of XRN2 (**Fig. 4a, S7a**). Notably, the unchanged nucleo-cytoplasmic distribution of a non-substrate miRNA, miR-21, upon XRN2 knockdown indicated towards the specificity of this entire sequence of events. However, it would require extensive work to know whether the miRNA releasing activities observed in these two fractions are contributed by different players or by the same factor, and whether they interact with XRN2 only or with other ‘miRNases’ as well.

### Comparative analyses of protein mediated release of miRNAs from Human AGO2 with miRNA release mediated by highly complementary target RNAs

Spanning over a decade, several reports have highlighted an alternative pathway for miRNA degradation, which primarily relies on release or tailing-and-trimming/ trimming-mediated destabilization of miRNAs from miRISC upon binding to target RNAs that possess extensive complementarity with the miRNAs^14,16,32,33^. Accordingly, *in vitro*-reconstituted AGO2-guide RNA complexes have been demonstrated to unload the guide RNA strand upon incubation with high molar excess of perfectly complementary target RNA sequences^32^. We therefore sought to perform a comparative study between the release of the AGO2-bound miRNAs mediated by the extensively complementary target RNAs, and the above-mentioned total cell lysate devoid of any nucleic acid due to treatment with micrococcal nuclease. We began by monitoring the release of let-7a from endogenous AGO2 in the presence of increasing concentration of an *in vitro*-transcribed target RNA, perfectly complementary to let-7a **(Fig. 5a-d**, **S7d**). We observed release of around 90% AGO2-bound let-7a upon incubation with ~150 folds molar excess of the target RNA at 37°C for 30 minutes **(Fig. 5d)**. Interestingly, very little or no miRNA was released from AGO2, when the reaction was performed with equimolar concentration of target RNA **(Fig. 5c).** Contrastingly, ~80% of endogenous AGO2-bound let-7a was released upon incubation with total lysate (75 μg), where amount of the lysate was in equivalence with that of the immunoprecipitated AGO2 employed in the ‘miRNA release assay’ **(Fig. 5c, 5e)**. Notably, with ~5% immunoprecipitation efficiency, the immunoprecipitated AGO2 employed in each miRNA release assay reaction was equivalent to ~75 μg of total lysate **(Fig. S7e)**, and from which ~20 femtomoles of let-7a could be extracted **(Fig. S7f)**. Upon incubation with 150 μg of total lysate, the release of let-7a was further facilitated to the extent of ~90% **(Fig. 5e)**. Similarly, let-7d, which was only 60% released upon treatment with 75 μg total lysate, showed a greater magnitude of release upon treatment with higher concentrations of total lysate **(Fig. 5e)**. These results clearly demonstrated that the release of miRNA from miRISC that could be achieved at a high molar excess of target RNA, could also be achieved with total protein lysate, but at a physiologically relevant concentration. Of note, in the aforementioned target-mediated release assay, we also measured the magnitude of release for other members of the let-7 family, for whom the employed let-7a-target RNA was only partially complementary. Indeed, the incompletely complementary target failed to efficiently release let-7i (4 mismatches) or let-7d (2 mismatches) from the endogenous AGO2, even when it was present at 150-fold molar excess **(Fig. 5d, S7g)**. Together, our data suggested that protein-mediated miRNA release from miRISC is likely to be an important mechanism for miRNA metabolism in humans.

**Figure 5.**
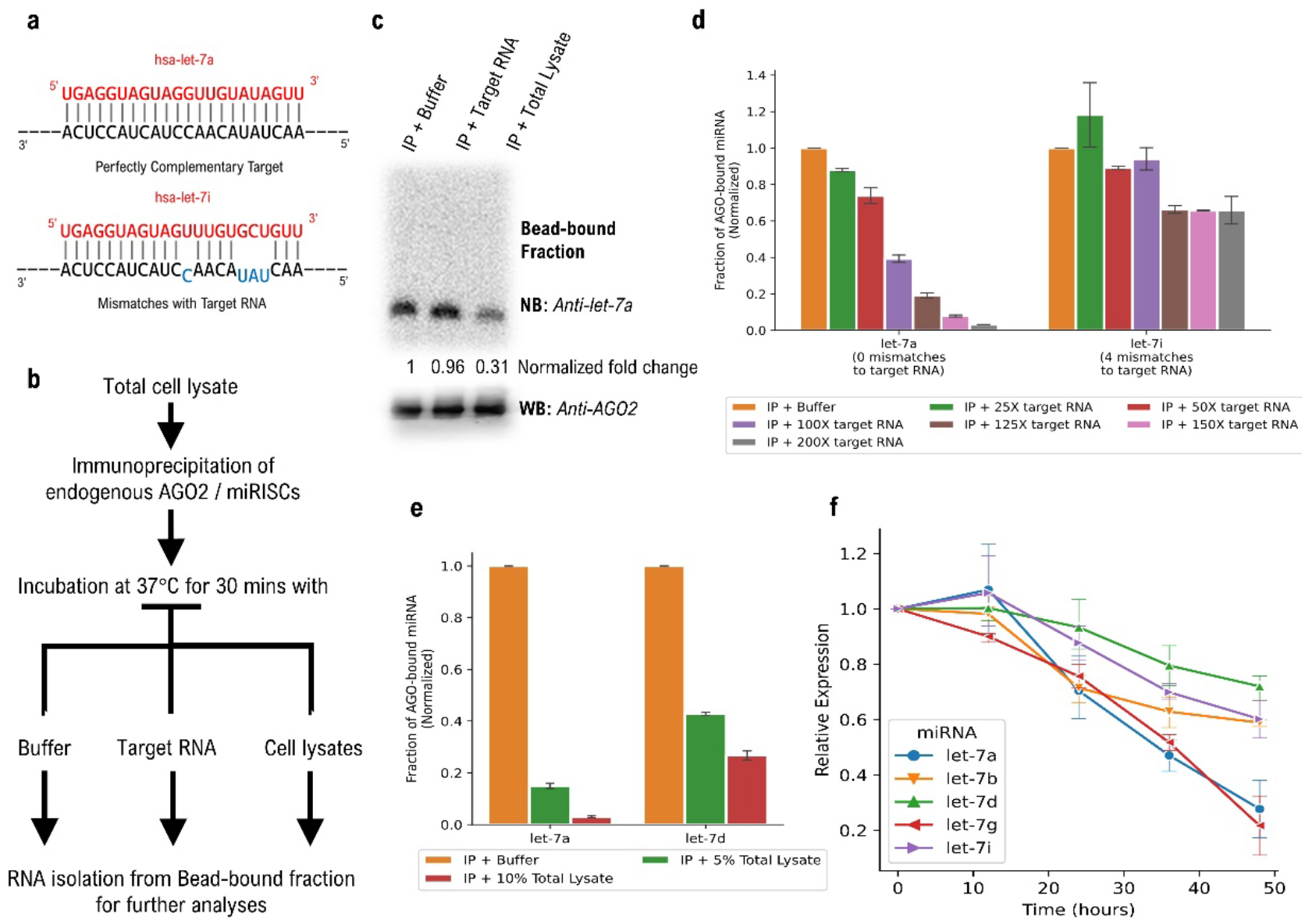
Nucleic acid depleted total lysate releases let-7 miRNAs from miRISC more efficaciously than completely complementary RNA. **a.** Schematic of *in vitro*-transcribed target RNA perfectly and imperfectly paired to let-7a and let-7i, respectively. **b.** Schematic representation of miRNA release assay to check the release of let-7 miRNAs from endogenous AGO/ miRISC mediated by complementary target RNAs or total cell lysate. **c.** Release of let-7a miRNA from immunoprecipitated AGO2 upon treatment with equimolar concentration of perfectly complementary target RNA or an amount of total cell lysate in equivalence with the employed AGO2. Post treatment, one half of the bead-bound AGO2 fractions were subjected to northern probing (top) and the other halves were subjected to anti-AGO2 western blotting acting as loading controls (bottom). **d.** TaqMan analyses (n = 4; mean ± SEM) reveal differences in the magnitude of release of two let-7 miRNAs from immunoprecipitated AGO2 upon treatment with increasing concentrations of a target RNA (perfectly complementary to let-7a, but with 4 mismatches for let-7i). **e.** TaqMan quantification (n = 4; mean ± SEM) of AGO2-associated let-7a and let-7d (re-recovered bead-bound fraction) upon treatment with different concentrations of total cell lysates. Let-7a is more amenable to release than let-7d. **f.** Total RNA was isolated from cells harvested at 0, 12, 24, 36 and 48 hrs following α-amanitin (10 μg/ml) treatment, and TaqMan quantification (n = 4; mean ± SEM) performed for the indicated let-7 members that reveal differences in their relative stabilities.

Interestingly, we observed that let-7b, let-7d and let-7i, unlike other let-7 family members, were not released to a substantial amount from the endogenous AGO2 upon exposure to the total lysate (Figure 4E). This could possibly be due to the following three reasons: (i) their release is partially dependent on target RNAs, (ii) the factor(s) responsible for their release from the specific miRISCs were partially inactive in the employed total lysate due to extraction conditions, or (iii) the composition of the respective miRISCs housing let-7b, let-7d and let-7i were such that they were resistant to the releasing activity, which might confer a greater stability and longer half-life to them. Indeed, by inhibiting RNA Pol II with α-amanitin, we could examine their relative stabilities, and they appeared to be more stable than the other members of the let-7 family **(Fig. 5f)**. This data suggested that the composition of a specific miRISC or a specific miRNP structure formed by a set of proteins and a given miRNA, could override a general turnover mechanism, and thus might play an important role in determining the functionality of the given miRNA.

### Analysis of clinical cancer transcriptomics data reveals the widespread impact of *XRN2* on miRNA homeostasis and cellular pathways

As previous reports have implicated ZSWIM8 Ubiquitin ligase and Tudor-SN (*SND1*) as two crucial regulators of miRNA abundance in mammalian cells^13,19,20^, we became interested to learn the relative contribution of these candidates towards negative regulation of the expression levels of the let-7 family members in a set of patients, where XRN2-mediated regulation of let-7 family members is favoured. In this direction, we compared the fold changes in the mean levels of let-7 family miRNAs between the low and high expression categories of *XRN2, SND1* and *ZSWIM8* mRNAs in the ‘representative subset’ for all three TCGA datasets defined earlier. In case of TCGA-HPCC subset, let-7a, −7c, −7d, −7f, −7g and miR-98 showed a stronger inverse relationship with *XRN2* mRNA than with those of the other two enzymes, which is further borne out by their increased accumulation in case of low *XRN2* mRNA levels than for the other enzymes considered **(Fig. 6a, 6d)**. However, in this subset, the expression of *SND1* mRNA also possessed an appreciable inverse relationship with some of the let-7 family members. Additionally, we found that the levels of a liver-specific miRNA, miR-122^2^, were also negatively affected upon increased expression of all three candidates, though maximally for *XRN2* **(Fig. 6a)**. Further we wanted to check the global picture of the inverse relationships of *XRN2, SND1*, and *ZSWIM8* mRNA levels with different let-7 family members. Accordingly, we repeated the comparison using the entire dataset. In the case of TCGA-HPCC, we observed a prominent and comparable negative effect of both *SND1* and *XRN2* mRNA expression on the levels of many let-7 family members **(Fig. S8a)**. However, high *ZSWIM8* mRNA expression appeared to have a much weaker effect on these miRNAs than the other two candidates. miR-122 was most negatively affected by XRN2 mRNA levels **(Fig. S8a)**. We also made observations from both the ‘representative subsets’ and the complete datasets for TCGA-GBM and TCGA-LUAD, employing the same method of analysis as for TCGA-HPCC **(Fig. 6b, 6c)**. All three candidate enzymes had comparable contributions to the negative regulation of let-7 family members in the ‘representative subset’ for TCGA-GBM, while in the whole dataset, *ZSWIM8* appeared to be the most prominent candidate, followed by *SND1* and *XRN2* **(Fig. S8c)**. This is consistent with previous reports, which suggested that TDMD might be more important phenomenon in the context of neuronal cells^16,33^. Notably, high mRNA levels of all three enzymes had a negative effect on the level of miR-124, a neuron-specific miRNA^2^, in the whole dataset for TCGA-GBM **(Fig. S8c)**. Also, in the complete dataset for TCGA-LUAD, it was *ZSWIM8* mRNA that had the strongest negative relationship with let-7 family miRNAs, followed by *SND1* and *XRN2* mRNA **(Fig. S8b)**. Although, in case of clinically important miR-29a^31^, *XRN2* was the only candidate to have an appreciable negative relationship in this complete dataset **(Fig. S8b)**. The above observations suggested that all three candidates have roles in regulating levels of let-7 family miRNAs, and their relative contribution towards maintaining let-7 levels vary considerably, not just in different tissues, but also in different cohorts of patients. XRN2 appears to have a markedly important role in regulating many let-7 family members, especially in hepatocellular carcinoma, and is a contributor to miRNA homeostasis in multiple cancer types, alongside other candidates implicated in miRNA turnover.

**Figure 6.**
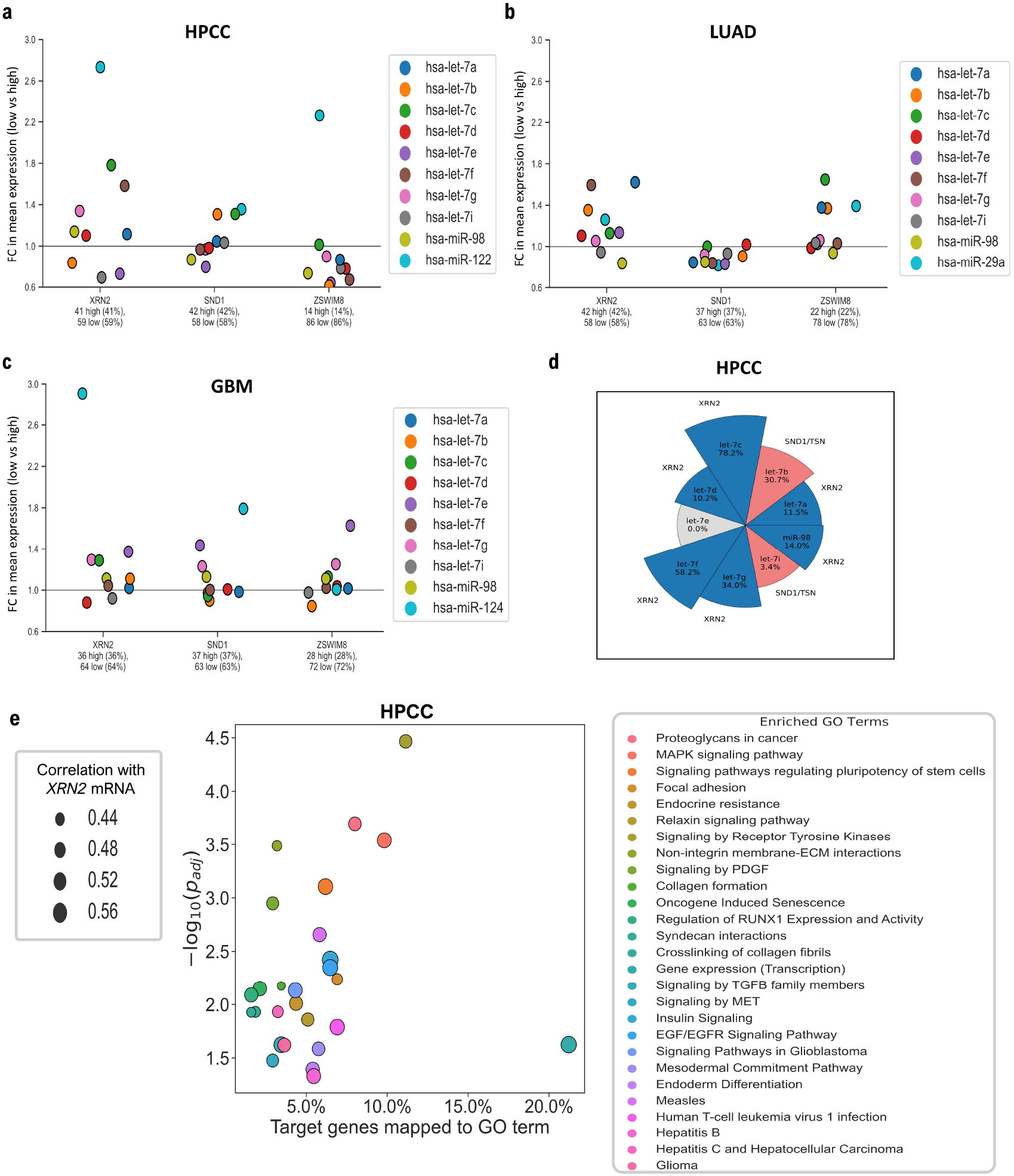
Widespread impact of XRN2-mediated regulation of human let-7 miRNAs on various cellular processes they are implicated in. **a-c.** Comparisons of the mean fold changes in expression of different let-7 family members and one tissue-specific miRNA between low and high expression groups for *XRN2, SND1* (Tudor-SN), and *ZSWIM8* mRNAs in the ‘representative subsets’ of the TCGA-HPCC (**a**), TCGA-LUAD (**b**), TCGA-GBM (**c**) datasets. Low *XRN2* mRNA levels are associated with different degrees of accumulation for the indicated miRNAs in the different subsets (eg. in TCGA-HPCC subset, low *XRN2* mRNA levels are associated with larger increases in let-7a, −7c, −7d, −7f, −7g, miR-98 and miR-122 levels in comparison to low expression of either *SND1* or *ZSWIM8* mRNAs). **d.** Maximum relative accumulation of let-7 sisters with low expression of *XRN2, SND1*, and *ZSWIM8* mRNAs from (**a**). Six out of nine let-7 members are maximally accumulated in case of low *XRN2* mRNA levels over low *SND1* or *ZSWIM8* mRNA levels. Amongst these, let-7a and let-7c display the most accumulation for low *XRN2* mRNA levels, while let-7b and let-7i are accumulated more for low *SND1* mRNA levels. let-7e is not appreciably accumulated for low expression of any of the three enzymes considered. **e.** Gene Ontology (GO) Term Enrichment analysis indicate the most enriched GO terms for the genes positively correlated with *XRN2* expression in the TCGA-HPCC dataset, include candidates from multiple signalling pathways, different types of cancers and viral infections, as well as pathways related to ECM metabolism. “Signalling by Receptor Tyrosine Kinases” is the most significantly enriched GO term (y-axis: significance of enrichment), while “Gene expression (Transcription)” of both the most abundant and enriched GO term (x-axis: fraction of analysed genes associated with a GO term) is also strongly correlated with *XRN2* mRNA expression (depicted marker radii on the left box are proportional to the mean Spearman correlation of all the GO-term associated genes with *XRN2* mRNA expression). This observation is consistent with many let-7 targets being transcription factors.

Furthermore, since, let-7 family members have been adequately characterised to be involved in regulating several important cellular pathways^24^, we went on to investigate if any other major cellular pathways, aside from those directly associated with cancers, were affected in relation to *XRN2* expression in the employed clinical tumour transcriptomics datasets. Gene Ontology (GO) terms are used to annotate genes with their functional roles in pathway databases such as the KEGG^34^ and Reactome databases^35^. Therefore, we performed GO term enrichment analysis on genes, which are predicted or validated targets of let-7 family members (as per human TargetScan 7.2). Among these genes, whose mRNAs were also positively correlated with *XRN2* mRNA in the relevant TCGA-HPCC dataset were selected for downstream analysis **(Fig. 6e)**. Given the roles of let-7 family members in regulating pluripotency, especially through the conserved LIN28B axis^24^, it was unsurprising to find that one of the enriched terms was related to regulation of stem cell pluripotency, and other terms related to differentiation. Similarly, due to the known roles of let-7 family members in regulation of genes involved in controlling the cell cycle like *CDK6, CDK8, CDC25A* and *CCND1*^24^, we also found terms related to cell cycle processes and mitotic checkpoints to be enriched. Enrichment of terms related to viral infections, especially of RNA viruses such as HTLV-1, measles, and Hepatitis C were also enriched. We additionally observed that regulation of cellular senescence through the p53 and Ras-MAPK/RTK signalling pathways^24^ were also enriched terms in the term enrichment analysis. Since, let-7 family members are also known to regulate the TGF-*β* receptor TGFBR1, as well as STAT3 and PI3K-AKT^24^, terms related to TGF-*β* and PDGF signalling were also enriched. Furthermore, it was interesting to observe that terms related to collagen biosynthesis, focal adhesion and syndecan interactions displayed enrichment as well. Notably, this could be because let-7a, let-7b and let-7g are known to regulate the expression of type 1 collagen genes^24^. Outcomes of this analysis suggest a strong involvement of both XRN2 and let-7 family members in regulation of cell proliferation and differentiation, as well as cell and extracellular matrix (ECM) homeostasis.

Very similar GO terms were enriched in the outcome of a similar analysis performed on the TCGA-GBM dataset **(Fig. S8d)**. The outcome of the GO term enrichment analysis performed on the TCGA-LUAD dataset was less intriguing, since the variety of enriched terms was much lower (unpublished observations). This could most probably be due to a higher sample heterogeneity in the TCGA-LUAD dataset as compared to the TCGA-HPCC and TCGA-GBM datasets, which might have allowed far fewer GO terms to reach the requisite significance threshold for representation. Notably, we found very similar GO terms enriched in the TCGA-HPCC and TCGA-GBM datasets, as well as cross-enrichment of hepatocellular carcinoma terms in GBM and glioma terms in HPCC. Probably, it indicates that the functional roles played by XRN2, and let-7 family miRNAs are preserved across different cell lineages, and further suggests far-reaching effects of XRN2 in various aspects of epithelial and glial development and homeostasis.

## Discussion

Our results demonstrate that in human cancer cells, turnover of let-7 family of miRNAs is a layered process involving multiple steps (Figure 7), and it acts in conjunction with other regulatory forces to govern the functionality of miRNAs. We furnish the role of XRN2 as a critical regulator of miRNA abundance with a prominent impact on the physiology of cancer cells, especially their tumorigenicity. Our findings suggest that the putative ‘miRNA release factor’ and XRN2-mediated miRNA decay might together constitute an important pathway to regulate the stability and function of let-7 members.

**Figure 7.**
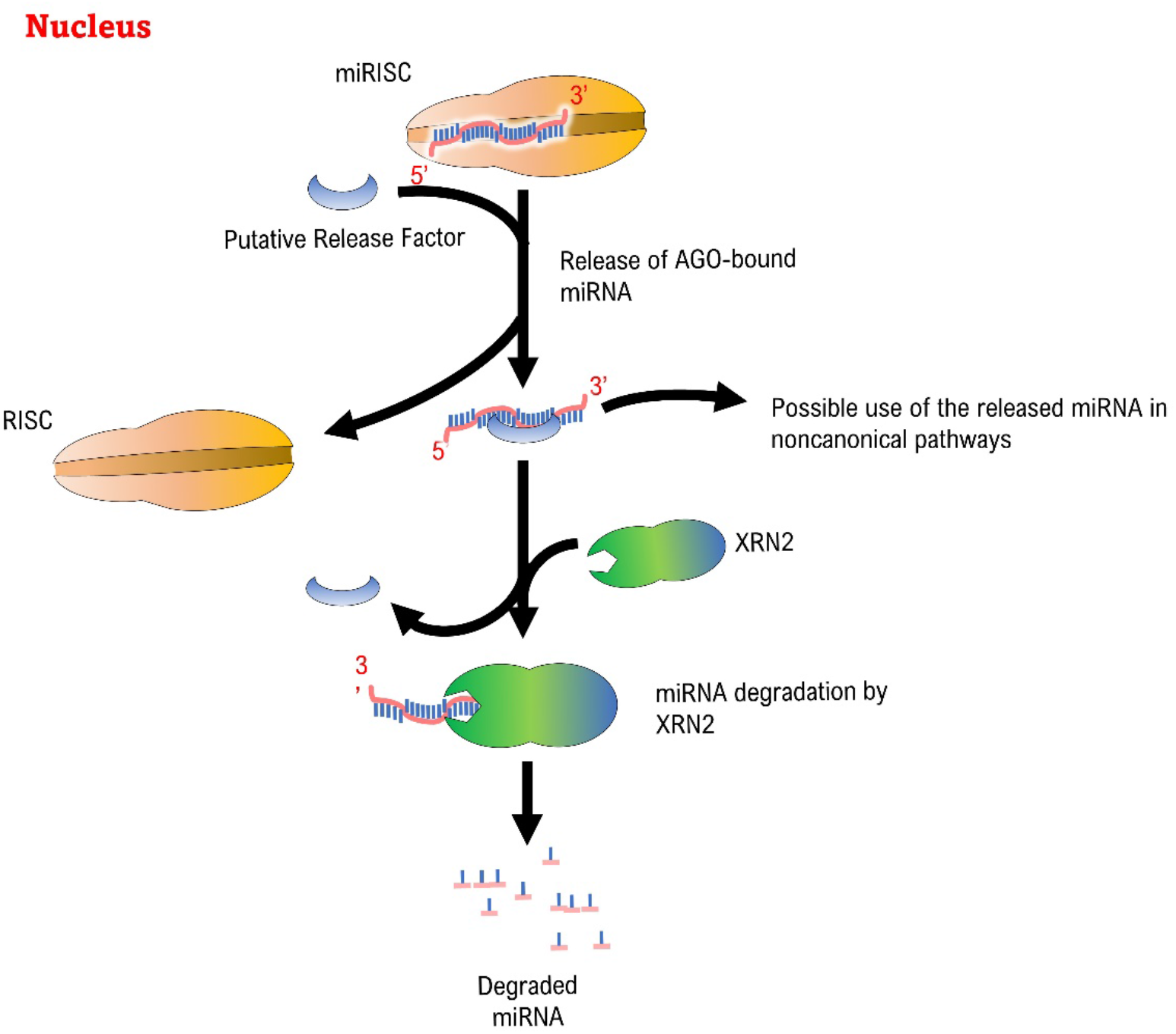
A putative model depicting XRN2-mediated miRNA turnover pathway. A two-step miRNA turnover model, where XRN2 acts on the miRNA upon its release from the AGO/miRISC by a yet unknown ‘miRNA release factor’.

A previous study reported that XRN2 binds pre-miR-10a in a DICER-independent manner and accelerates the maturation of miR-10a in human lung cancer cells^25^, but did not provide insights on the members of let-7 family. Another study demonstrated reduced degradation of a synthetic miRNA by an XRN2-depleted lysate but did not augment the observation through *in vivo* evidence^36^. Here, we demonstrate that in human cancer cell lines of lung, hepatic, and neuronal origin, XRN2 depletion led to the accumulation of a number of mature let-7 family members without any effect on their pri- or pre-miRNAs. Furthermore, many other known tumor suppressor family members also showed accumulation of their mature forms. However, the factor(s) determining the specificity of XRN2 for its substrate miRNAs need to be determined. Since, XRN2-mediated miRNA degradation happens downstream of a miRNA release process, it is very likely that the specificity of interaction between this putative ‘release factor’ and different miRISC complexes, as well as with XRN2 would determine the final outcome. Thus, identification of the miRNA release factor and understanding its interaction with XRN2 hold key to further unravel the question of specificity.

We have also presented that XRN2 depletion in the chosen cancer cells led to a significant decrease in the abundance of several oncogenic mRNAs that are validated targets of let-7 and other accumulated tumor suppressor miRNAs. Since, most of those oncogenic mRNAs host multiple miRNA binding sites, understanding of the relative contribution towards regulation of those targets by the different tumor suppressor miRNAs would require detailed analyses that will keep multiple determinants of miRNA-mediated regulation under consideration (eg. relative concentration of the different miRNAs, number of same/different miRNA binding sites and relative distance between those sites, target site architecture, expression of miRNA specific factors that facilitate/antagonize miRISC function etc.). Nevertheless, it was clear that XRN2-mediated regulation of miRNAs indeed plays a vital role in shaping the transcriptomic landscape of lung cancer cells. The pathophysiological significance of our findings was reinforced not only by the xenograft experiments using nude mice, but also through the analyses of public tumour transcriptomics data from LUAD, HPCC, and GBM patients, especially considering the heterogeneity of the patient cohorts. Those analyses demonstrated an appreciable inverse relationship between the levels of certain let-7 family members and *XRN2* mRNA in the clinical tumour samples, which corroborated with our cell line-based observations. They also revealed that let-7 target genes positively correlating with *XRN2* mRNA expression are involved in multiple signalling pathways regulating cell proliferation and differentiation, as well as in the biogenesis and maintenance of extracellular matrix in both epithelial and neuroglial cell lineages. This partially reveals why elevated *XRN2* has previously been associated with worse survival outcomes in lung cancer patients, and also accentuates prior observations on overexpressed *XRN2* mRNA in spontaneously occurring lung tumours in mice and human tumour microarray data analysed through genome-wide association studies^22^. Apart from let-7 members, XRN2 depletion also affected the levels of certain tissue-specific miRNAs, like miR-122 in hepatocellular carcinoma cells, and miR-124 in case of glioblastoma cells, which are known to have important functions in the developmental processes of liver and neuronal tissues, respectively, and whose dysregulations have been linked to multiple disease states^2^.

Our observations from the analyses of TCGA-LUAD, TCGA-HPCC and TCGA-GBM datasets indeed revealed the association of high *XRN2* mRNA levels with worse survival in LUAD, HPCC and GBM patients, respectively. Our motivation towards performing bioinformatic analyses employing the TCGA datasets was because of how they encompass samples from a large and diverse patient population for each cancer type and the corresponding datasets, while ensuring that all sequencing data acquisition and processing have been performed using uniform procedures. We expect that our observations, especially considering the large and heterogenous clinical population represented in the TCGA datasets, would advocate for wider applicability of the molecular signatures of XRN2 in determining cancer severity than what could be suggested from smaller or more limited datasets. With the revelation of a clear inverse relationship between the levels of let-7 members and *XRN2* expression, we found that for two other important factors implicated in miRNA turnover, such an inverse relationship between the let-7 family members and those concerned mRNAs (*SND1* and *ZSWIM8*) was comparatively weaker in patient-derived samples favouring XRN2-mediated let-7 regulation. In the larger complete datasets, we found that XRN2 is a critical player in maintenance of let-7 miRNA levels in HPCC as well as in GBM and LUAD, alongside the other two critical players implicated in miRNA turnover (Tudor-SN and ZSWIM8). These observations indeed remind us of the fact that complexity of cancer is often an outcome of perturbations of multiple regulators acting in parallel. Therapeutic approaches relying on targeting miRNAs, even from the same family, would be greatly benefited by a more thorough understanding of the landscape of miRNA turnover, potentially allowing the design of drug formulations that can target multiple such pathways at the same time.

It has been previously described that exogenously introduced target RNAs having extensive complementarity to the cognate miRNAs can lead to target RNA–directed tailing and trimming of endogenous miRNAs, and influence their stability and function^14^. Structural studies have demonstrated that the 5’-end of a miRNA remains anchored in a pocket in the MID domain of AGO, and the 3’-end is bound by the PAZ domain. The miRNA remains largely buried in a channel between the two lobes of AGO, except the bases of the miRNA-seed region, which remain partially exposed to the surface of the host protein. However, extensive base pairing of AGO2-bound miRNA with target RNAs disrupts the 3’-end anchoring and renders the 3’-end accessible for enzymatic activity^37^. Also, specific AGO-bound miRNAs that have their 3’-ends exposed, have been demonstrated to undergo an oligo-tailing, followed by degradation by DIS3L2^18^. Since, for most cellular AGO-bound miRNAs, their 3’-ends are protected by the AGO proteins, unless they encounter extensively complementary targets, it is improbable that DIS3L2 mediated miRNA turnover pathway could be the foremost channel for miRNA turnover inside the cell^18^. Our studies provide evidence that XRN2-mediated degradation of miRNAs is dependent on an upstream event of ‘miRNA release’ from miRISCs, carried out by an yet unidentified proteinaceous factor(s). Moreover, our *in vitro* results indicate that protein factor-mediated release of AGO-bound let-7 appears to be far more efficacious compared to complementary target RNA-mediated mechanism^32^. Additionally, this two-step turnover mechanism has the potential to prevent immediate degradation of an AGO-released miRNA, which might enable shuttling of some specific candidates to the yet uncharacterized non-canonical pathways that utilize mature miRNAs without any association with known components of miRNA metabolism like AGO, GW proteins, etc.^38,39^, whereas most of the released miRNAs might get delivered to the bona fide ‘miRNases’ for degradation.

Interestingly, the recently reported ‘miRNase’-Tudor-SN co-immunoprecipitates with RISC subunits and appears to be a component of miRISC^13^. The intracellular signalling events marking an AGO-residing miRNA for termination of its function remain largely unknown, and it will be important to know the responsible factors and the mechanism that allows such miRNAs to travel from the grasp of AGO to Tudor-SN for degradation.

Downregulation of endogenous miRNAs has been shown to be induced by certain viral RNAs^40–43^, as well as a few genome-encoded transcripts^16^, which bear extensive complementarity to specific miRNAs. Recently, highly complementary target interactions were demonstrated to induce TDMD, independently of tailing and trimming events, by causing destabilization of AGO proteins through the ZSWIM8 ubiquitin ligase^19,20^. It has been proposed that ZSWIM8 directly recognizes the conformational changes induced in AGO upon extensive base-pairing of the resident-miRNA (miR-7) with a highly complementary target (Cyrano lncRNA), but the mechanism for this recognition remains to be understood. Since it is well known that 3’-tailing often acts as an important signal for downstream processes^14,33,44^, in its absence, it is quite possible that there might be yet unknown factor(s) that would facilitate recognition of the conformational changes elicited upon highly complementary miRNA-target interaction by ZSWIM8 to mediate TDMD. Additionally, the nucleases participating in the degradation of miRNAs, post their release from AGO undergoing proteolysis, needs to be identified. It is also noteworthy that the half-lives of AGO proteins are longer than most of the miRNAs they host^5,6^, suggesting that miRNA release and degradation through destabilization of AGO by ZSWIM8 and related factors is unlikely to be the major pathway for miRNA turnover. Intriguingly, a highly complementary viral RNA element-mediated degradation of a host miRNA may be considered an efficient viral strategy to destabilize the host system. However, turnover of a miRNA due to highly complementary interaction with the cognate endogenous target, mediated by destabilization of the AGO itself, would certainly impose an energy burden on the cellular system. Importantly, such unusual endogenous targets are not plentiful, and thus, TDMD through destabilization of AGO may have evolved as a regulatory measure under special circumstances for a specific set of miRNAs with exclusive functions. Contrary to TDMD, pairing of target mRNA with its cognate miRNA has been shown to exert a protective effect on miRNAs of specific families from cellular degradation in *C. elegans*^9^. Furthermore, target mediated protection of a few miRNAs was also reported to be crucial for controlling miRNA levels and their subcellular localization in human cells^45^. Also, in human cells, biogenesis of specific miRNAs was stimulated upon binding to their cognate target mRNAs^46^. Collectively, these reports indicate that probably there is no single principle and mechanism of turnover that determine the stability or vulnerability of miRNAs in different tissues and organisms. Crucially, while TDMD has been suggested to be an important mechanism for miRNA homeostasis in neuronal cells^33^, highly complementary let-7 targets suitable for TDMD are not yet known. Given our observations concerning *ZSWIM8* mRNA levels being prominently associated with a negative effect on let-7 levels in GBM and LUAD, ZSWIM8 might be involved in additional pathways to exert its regulatory effect in patient samples of these cancer types, and further exploration of those regulatory roles would be critical for better understanding of disease states.

Alongside their well-studied roles as tumour suppressors, let-7 family miRNAs are also important for mounting immune responses in vertebrates upon exposure to pathogens^47^. It is known to repress both the pro-inflammatory cytokine interleukin 6 (IL-6) and the anti-inflammatory cytokine IL-10 during *Salmonella* infection in murine phagocytic RAW264.7 cells, and thereby regulate early and late immune responses^47^. Interestingly, XRN2 depletion in RAW264.7 cells led to the upregulation of certain let-7 family members, and that in turn affected the levels of interleukins (unpublished observations), and this clearly makes XRN2 a potential candidate for modulation of inflammatory responses during infection. Notably, many of the patients infected by SARS-CoV-2 have been reported to show uncontrolled elevation in cytokines with persistent acute inflammation. Failure of the cellular homeostatic machinery to appropriately downregulate the ‘cytokine-storms’ leads to a vicious cycle of accumulating inflammation and subsequent lethality of the condition^48^. Since, let-7 has been shown to be capable of targeting multiple interleukins, its regulation through XRN2 might play a vital role in such critical situations. Moreover, IL-6 is also implicated in regulating various hypothalamic functions and contributes to the activation of hypothalamo-hypophyseal-pituitary axis, and thus to overall metabolism in vertebrates^49,50^. Supported by our cell culture-based results employing neuronal cancer cells and patient samples of neuronal tissues, it is possible that in the aforementioned neuro-endocrine tissues, let-7 plays a modulatory role on IL-6, and that in turn gets regulated by XRN2. Overall, these evidence and suggestions indicate that the regulation of let-7 through XRN2-mediated degradation might be conserved across the eukaryotic systems, and have widespread implications ranging from cancer physiology to host responses during infection. Further exploration of the functions of XRN2 and other components of the XRN2-mediated miRNA turnover pathway could shed new light on how these crucial regulators of the transcriptome themselves get regulated, and thus, might also provide opportunities for the discovery of new diagnostics and therapeutics.

## Author Contributions

SC designed the research and experiments. RN and TC performed the experiments. Together, all the authors analyzed the results and wrote the manuscript.

## Acknowledgments

We thank Pradipta Kundu for technical assistance with cell culture, and K. Somasundaram, Samarjit Jana for help with animal experiments. Results presented here are in part based on analyses performed with data generated by the TCGA Research Network: https://www.cancer.gov/tcga, specifically the TCGA-HPCC, TCGA-GBM, and TCGA-LUAD datasets. We are grateful to the specimen donors and research groups, who made the TCGA initiative possible. This work has been partly supported by the SERB/ DST (Govt. of India). RN received CSIR-Syama Prasad Mookerjee Fellowship (Govt. of India), and TC was supported by KVPY/ MHRD (Govt. of India).

