## Supplementary Information for "XRN2-mediated regulation of tumor suppressor microRNAs is of critical pathophysiological significance in Humans"

**This file includes:**

**Supplementary Text 1, 2**

**Supplementary Figures S1-S8 with their legends**

**Supplementary table 1**

**Materials and Methods**

**Supplementary Text 1**

**Efficient and specific depletion of human homologue of the worm ‘miRNase’ XRN-2 in human cell lines.**

In *C. elegans*, both XRN-1 and XRN-2 play the role of a ‘miRNase’^2,10^. In case of human cells, XRN1 has been implicated in the decay of miR-382 in HEK293 cells, whereas authors did not find any significant effect on the level of miR-382 upon depletion of XRN2^11^. They performed shRNA-mediated knockdown of *XRN1* and *XRN2* but did not demonstrate the knockdown efficiency at the protein level. Notably, depletion of XRN2 in A549 and H441 cells was shown to affect the maturation of miR-10a, leading to a decrease in the levels of mature miR-10a^12^. In this report, the authors mentioned a few downregulated miRNAs, and focused on the one getting most downregulated (miR-10a) but did not discuss the miRNAs that might have been upregulated. Additionally, in these studies, a substantial knockdown of XRN2 was not achieved^11,12^, which could be insufficient to exert an effect on all the pathways XRN2 is involved in. Therefore, prior to investigating the role of XRN2 in miRNA degradation in human cells, we optimized a rapid and efficient shRNA-mediated specific knockdown of *XRN2* using a lentivirus-based approach. We were able to achieve rapid (a timepoint of ~72 hr post transduction) and substantial knockdown of *XRN2* gene expression, with a knockdown efficiency of ~75% at the protein level in multiple cell lines (*e.g*., A549, MDA-MB-231, HEK293T; **Fig. S1**). Since, *XRN2* shRNA1 showed the best knockdown efficiency, it was used for all the subsequent experiments. To confirm that the shRNA is specifically targeting *XRN2*, and not *XRN1*, we checked the level of XRN1 protein in the XRN2-depleted cells. No effect on XRN1 level in the XRN2 knockdown cells indicated that shRNA was indeed specific for *XRN2* (**Fig. S1a**, middle panel). We also checked the level of AGO2 protein in the XRN2-depleted cells, which also remained unaffected with the knockdown of XRN2 in A549 cells (**Fig. S1a**, bottom panel). Notably, XRN2 is known to affect the overall RNA metabolism due to its role in the biogenesis of 5.8S rRNAs^14^. But, at our time-points, the levels of these rRNA remained unaffected upon depletion of XRN2 (**Fig. S1d**).

**Supplementary Text 2**

**let-7i shows limited or no regulation by XRN2.**

A perfect let-7a-target mRNA can also be a target for the other members of the let-7 family. But the magnitude of regulatory effect of a given member would be dependent on its abundance, coherence with the target expression, as well as its subcellular localization; thus, making a given target mRNA relatively more specific to a particular family member. We observed that let-7i showed very little or no changes in their levels upon XRN2 knockdown in the examined cell lines. And let-7i was also not negatively correlated with *XRN2 mRNA* in any of the TCGA-HPCC, TCGA-LUAD or TCGA-GBM datasets **(Fig. 2i-k)**. To confirm that the unaffected levels of let-7i were indeed manifested functionally, we checked the levels of known let-7i target mRNAs, namely, *KLK6* (let-7i target)^29^*,* and *PLK1*. As expected, levels of these mRNAs were not affected upon XRN2 depletion **(Fig. S3g-i)**.

**Supplementary Figures and Legends**

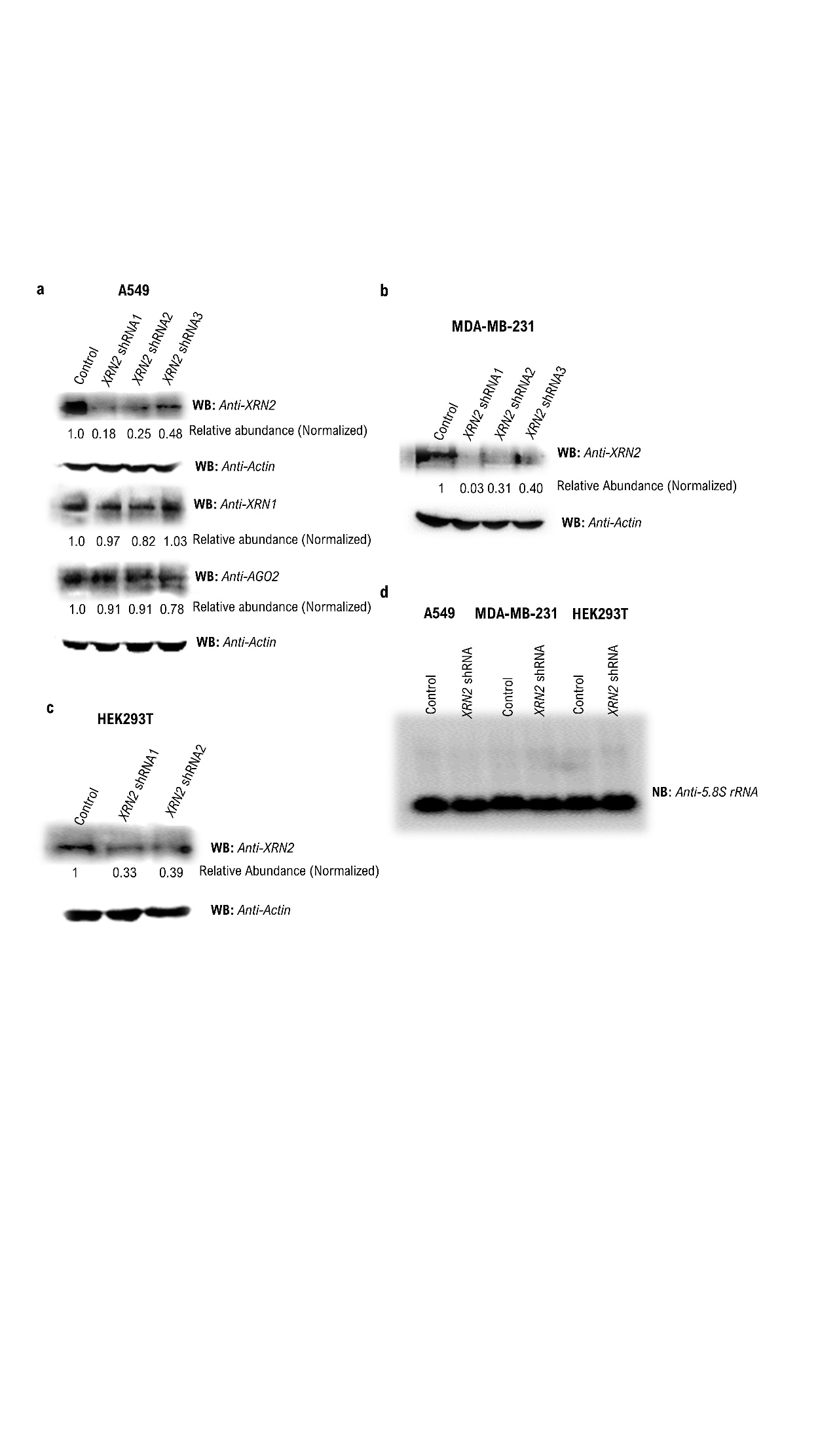

**Figure S1. Efficient and specific knockdown of XRN2 in different human cell lines.**

**a-c.** Efficiency of lentivirus-based shRNA mediated knockdown of XRN2 in different cell lines as indicated, was determined by western blot analyses using antibodies against XRN2. Unchanged levels of XRN1 and AGO2 (EIF2C2) proteins indicate the specificity of the knockdown.

**d.** Levels of 5.8s rRNA remain unchanged upon XRN2 knockdown.

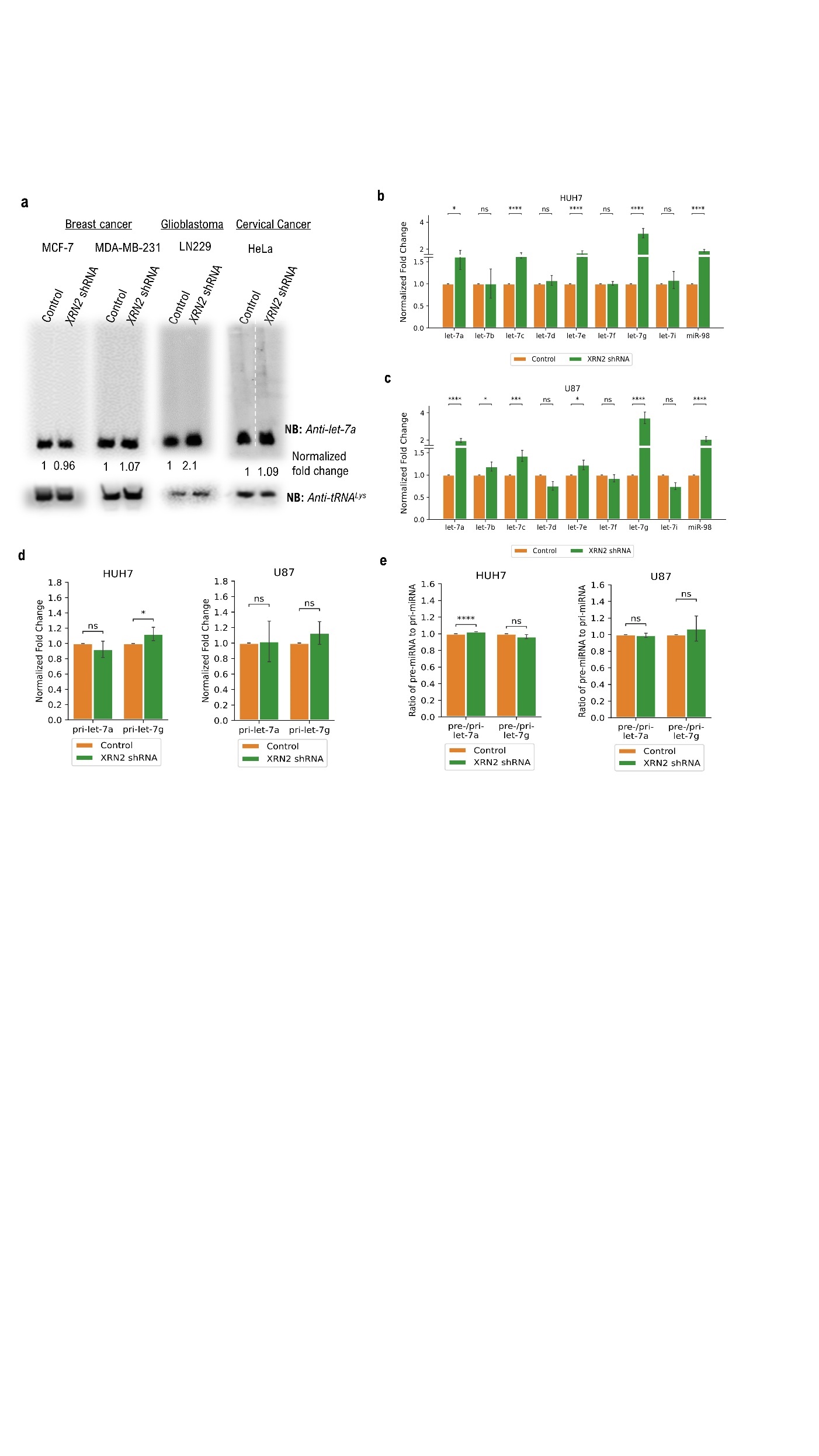

**Figure S2. Effect of XRN2 depletion on the members of let-7 family and their precursors in different cell lines.**

**a.** Effect of XRN2 depletion on levels of mature let-7a in MCF7, MDA-MB-231, LN229, and HeLa cell lines. Unrelated lanes were removed from the right-most panels.

**b.** Levels of mature miR-21 remain unaltered upon XRN2 depletion in A549 cells.

**c.** TaqMan analyses reveal increased abundance of the mature forms of different let-7 family members upon depletion of XRN2 in HUH7 (top panel) and U87-MG (bottom panel) cell lines. RNU48 RNA served as the normalization control.

**d.** RT–qPCR (n = 4; mean $\pm$ SEM) analyses reveal no significant change at the level of primary RNAs for the indicated miRNAs in HUH7 and U87 cells.

**e.** RT-qPCR (n = 4; mean $\pm$ SEM) analyses depict no significant change in the relative levels of pre-miRNA and pri-miRNA transcripts for indicated miRNAs in HUH7 and U87 cells.

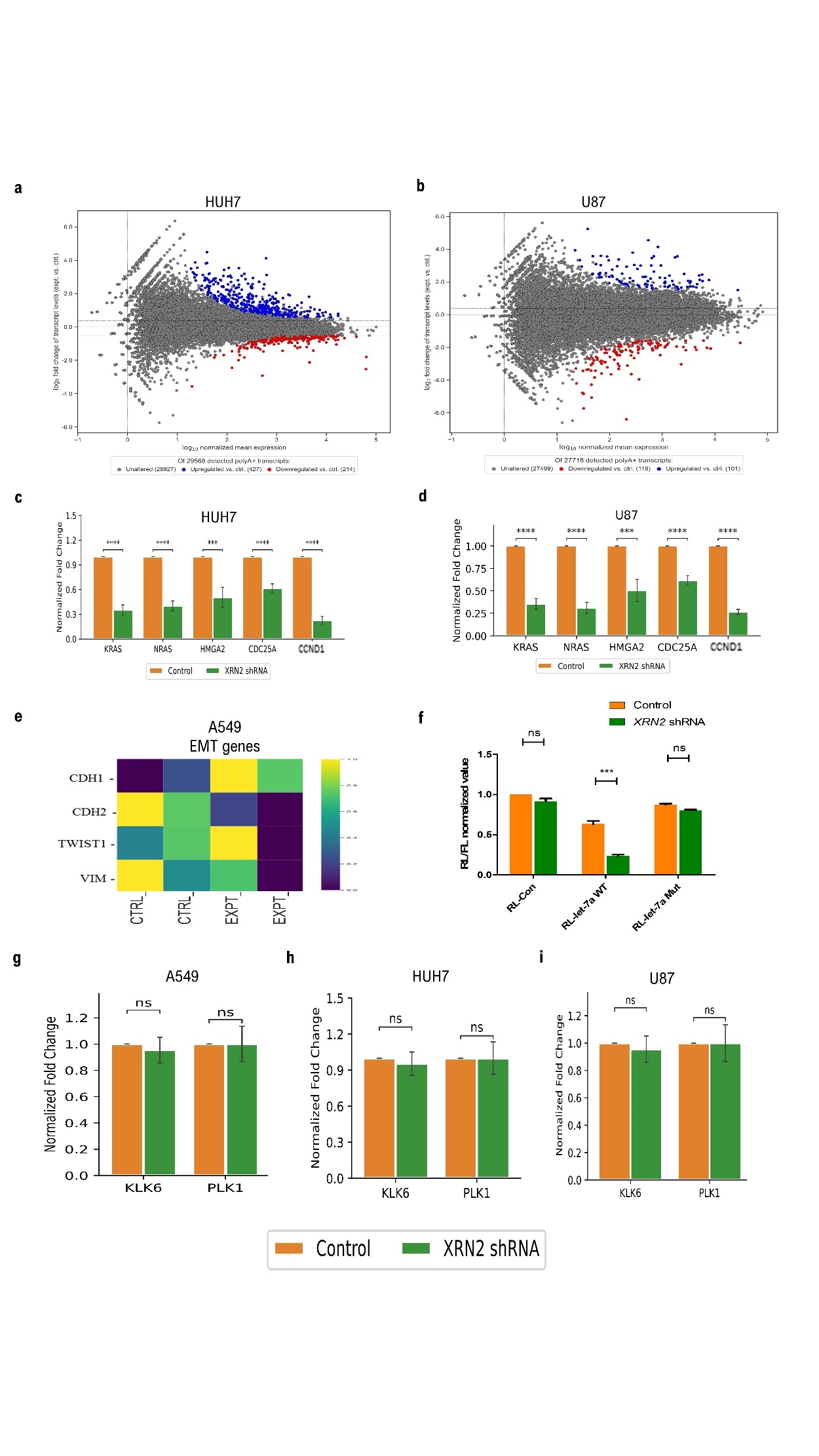

**Figure S3. Effects on endogenous and reporter mRNA levels upon XRN2 knockdown.**

**a.** MA/ Bland-Altman plot of differential expression of detected polyA+ transcripts for annotated genes in XRN2-depleted HUH7 cells. In red are genes whose transcripts are downregulated at least 0.7× with respect to the control samples (214/29568, 0.7%), and in blue are genes whose transcripts are upregulated at least 1.3× with respect to the control samples (427/29568, 1.4%).

**b.** MA/ Bland-Altman plot of differential expression of detected polyA+ transcripts for annotated genes in XRN2-depleted U87 cells. In red are genes whose transcripts are downregulated at least 0.7× with respect to the control samples (118/27718, 0.4%), and in blue are genes whose transcripts are upregulated at least 1.3× with respect to the control samples (101/27718, 0.3%).

**c, d.** RT–qPCR (n = 4; mean $\pm$ SEM) analyses reveal reduction in let-7 target mRNAs, as indicated, upon XRN2 depletion in HUH7 and U87 cells.

**e.** Relative expression levels (arbitrary units) of the EMT genes derived from the mRNA-seq data of control and XRN2 knockdown A549 samples.

**f.** Bar diagram depicting decreased Luc activity from cells transfected with Renilla luciferase construct hosting 3x let-7a target sites in 3’-UTR (**RL-let-7a WT**) upon XRN2 depletion, compared to cells transfected with control constructs (without let-7a target sites in the 3’-UTR, **RL-Con**) or constructs hosting mutations in the let-7a target sites (**RL-let-7a Mut**). These results indicate increased expression of let-7a upon XRN2 knockdown and sequence based interaction between let-7a and its target sites is key to the observed regulation.

**g-i.** mRNA targets of let-7i remain unaffected in XRN2-depleted A549, HUH7 and U87 cells.

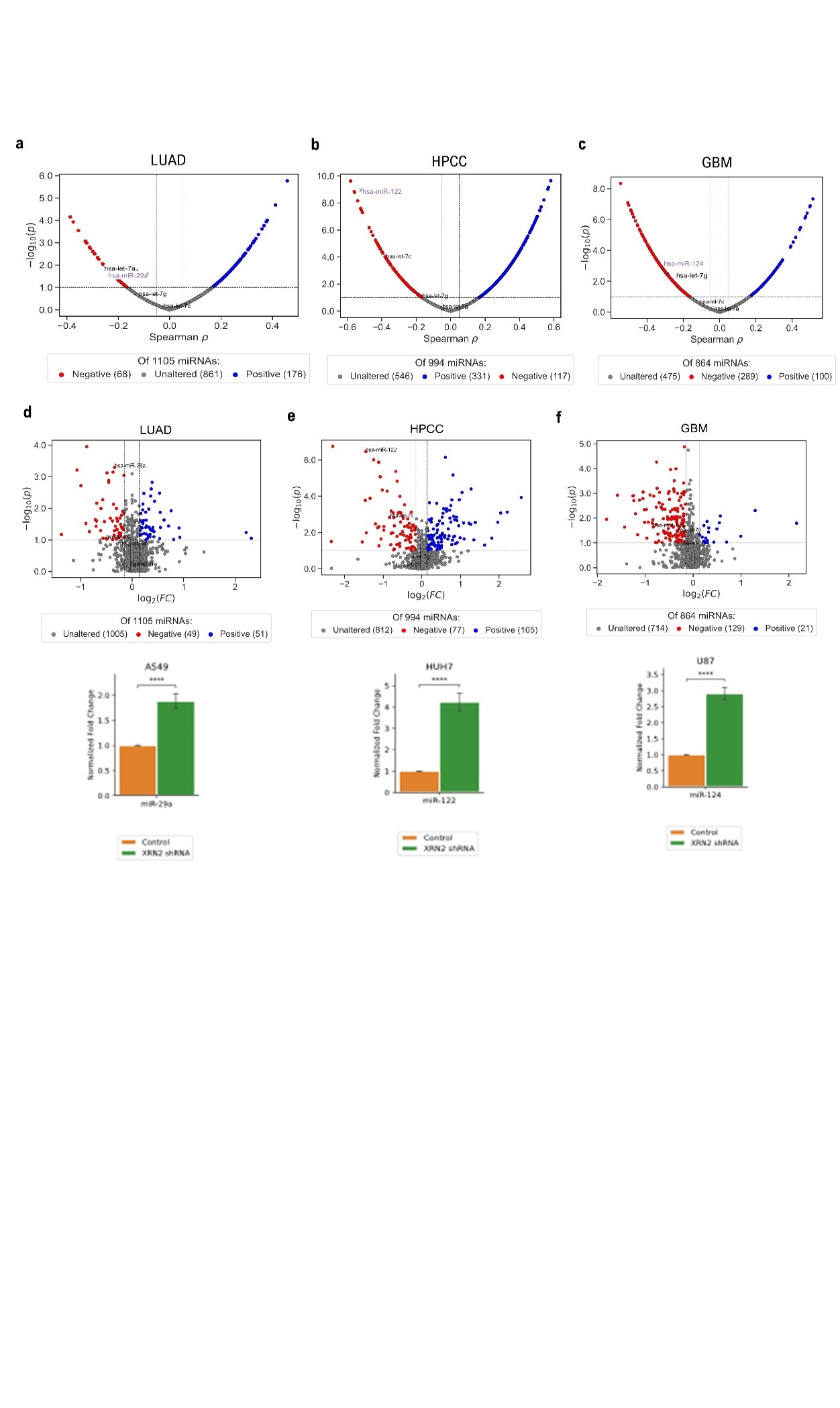

**Figure S4. Increased *XRN2* mRNA levels negatively influence the expression of a large number of miRNAs in patient (TCGA-LUAD, TCGA-HPCC, TCGA-GBM) datasets.**

**a-c.** Volcano plots of significance of fold change (y-axis: -log_10_(p)) vs. Spearman $\rho$correlation (x-axis: Spearman $\rho$) of all detected miRNAs with *XRN2* mRNA expression in the ‘representative subsets’ for TCGA-LUAD (**a**), TCGA-HPCC (**b**), and TCGA-GBM (**c**), and further confirms the inverse relationship of three let-7 sisters (let-7a, -7c, -7g) in the “representative subsets” for each cancer dataset, as well as that of the specific miRNAs miR-122 (HPCC), miR-124 (GBM) and miR-29a (LUAD) with *XRN2* mRNA. Correlation thresholds indicated are -0.05 and +0.05, and the significance threshold indicated is p = 0.1.

**d-f.** **Top Panels.** Volcano plots of significance of fold change (y-axis: -log10(p)) vs. mean fold change (x-axis: log2(FC)) of all detected miRNAs between high and low XRN2 mRNA expression groups in the ‘representative subsets’ for the TCGA-LUAD (**d**), TCGA-HPCC (**e**), TCGA-GBM (**f**) datasets, as depicted. With respect to XRN2 mRNA levels, the number of miRNAs detected to be significantly negatively altered in TCGA-HPCC (e) was 77, which included let-7c and the liver-specific miRNA miR-122, while 153 miRNAs were negatively altered in TCGA-GBM (f) including the neuron-specific miRNA miR-124, and among the 36 miRNAs negatively altered in TCGA-LUAD (d) was the clinically important miR-29a. **Bottom Panels.** TaqMan analyses of the levels of indicated mature miRNAs upon XRN2 depletion in three different cell lines, as depicted.

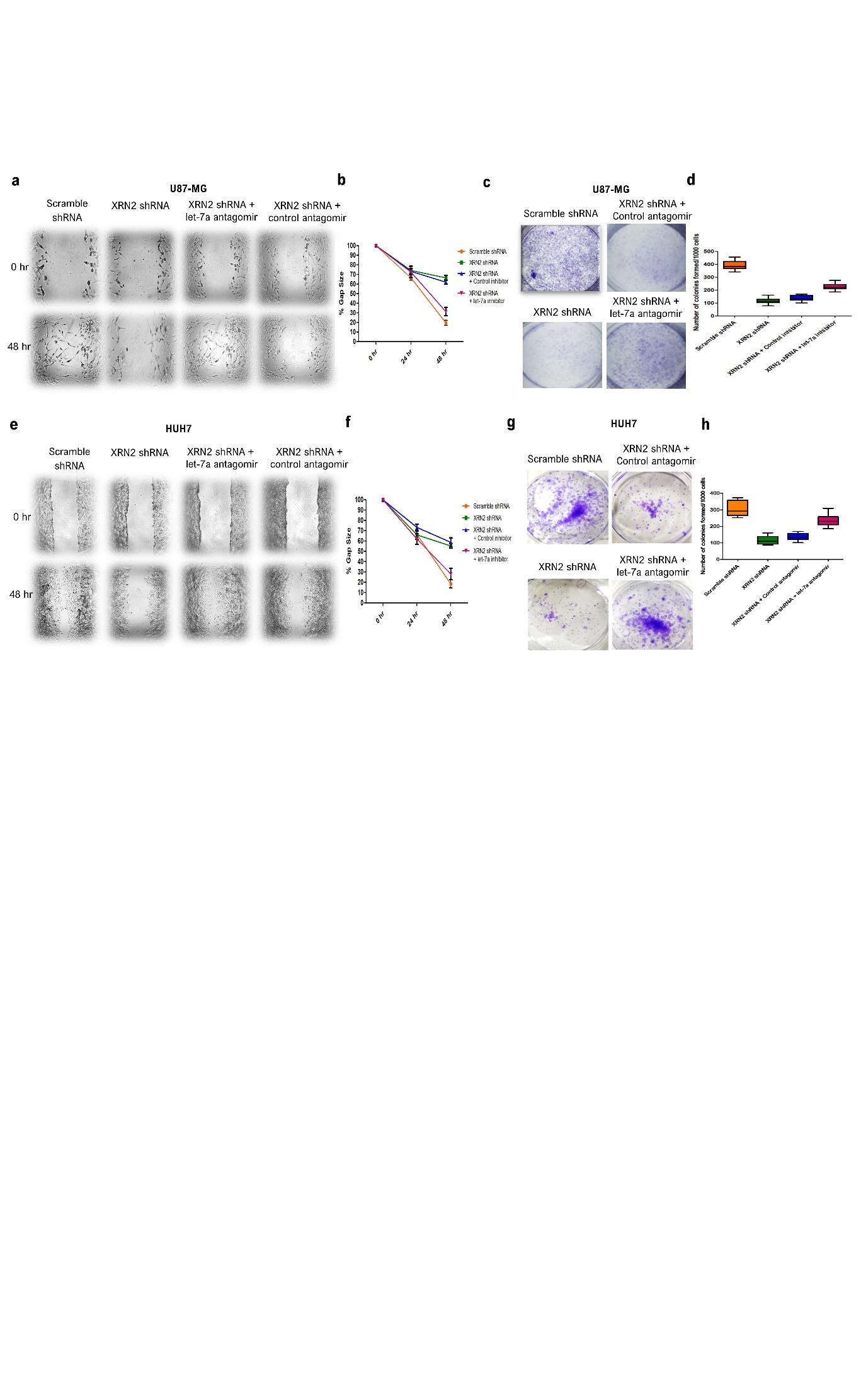

**Figure S5. XRN2 depletion affects cellular physiology of HUH7 and U87 cells.**

**a, b.** Wound-healing assay performed with U87-MG cells (n=4) transfected with indicated shRNAs depict reduced cell migration in cells transfected with XRN2 shRNA compared to scramble shRNA. Treatment with let-7a antagomir efficiently reverses decreased cell migration of XRN2-depleted U87-MG cells. Graphical representation of the wound-healing assay reveals significant differences in the gap-size between the indicated samples.

**c, d.** Clonogenic assay performed with U87-MG cells employing scramble shRNA and XRN2 shRNA (n=4) depicts decreased colony formation ability of XRN2 depleted cells. Corresponding box plot demonstrates the decrease in the number of colonies formed by XRN2-depleted cancer cells and reversal of the phenotype upon subjecting let-7a antagomir treatment to the XRN2-depleted cells.

**e, f.** Wound-healing assay performed with HUH7 cells (n=4) transfected with indicated shRNAs depict reduced cell migration in cells transfected with XRN2 shRNA compared to scramble shRNA. Treatment with let-7a antagomir efficiently reverses decreased cell migration of XRN2-depleted HUH7 cells. Graphical representation of the wound-healing assay reveals significant differences in the gap-size between the indicated samples.

**g, h.** Clonogenic assay performed with HUH7 cells employing scramble shRNA and XRN2 shRNA (n=4) depicts decreased colony formation ability of XRN2 depleted cells. Corresponding box plot demonstrates the decrease in the number of colonies formed by XRN2-depleted cancer cells and reversal of the phenotype upon subjecting let-7a antagomir treatment to the XRN2-depleted cells.

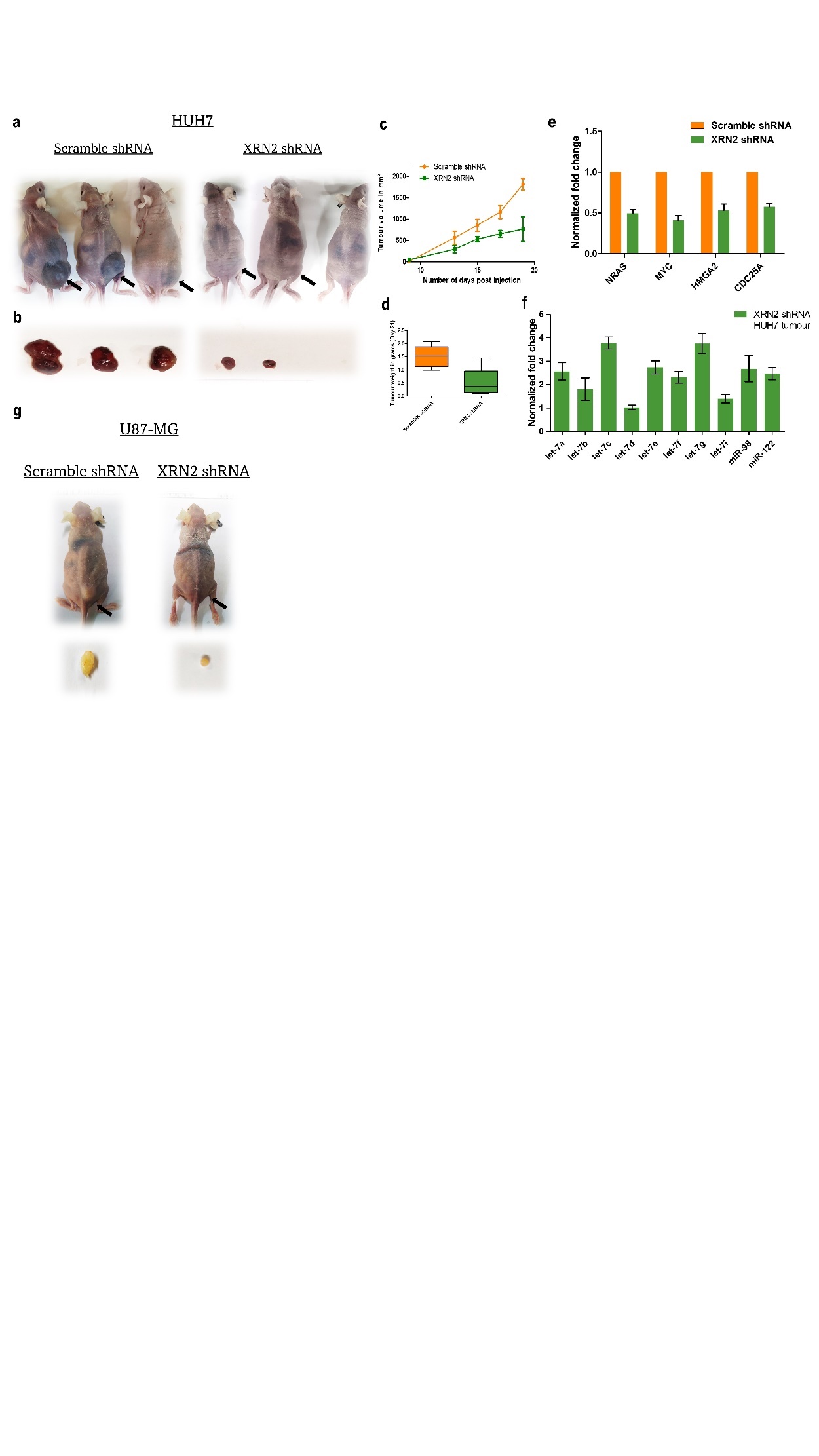

**Figure S6. XRN2 depletion affects physiology of cancer cells in mouse xenograft	experiments.**

**a, b.** Representative images of mice injected subcutaneously with either scramble shRNA or XRN2 shRNA transfected HUH7 cells after 21 days of injection. (**c, d**) Graph representing the tumor volume (mm^3^) and tumor weight in grams (gm) for scramble shRNA and XRN2 shRNA transfected HUH7 tumors after 21 days of injection.

**e.** RT-qPCR analyses reveal downregulation of expression of let-7 target mRNAs in XRN2-depleted HUH7 tumors compared to control tumors. RT-qPCR (n = 5; mean ± SEM) results were normalized against ATP5G mRNA.

**f.** TaqMan analyses reveal increased abundance of the mature forms of different let-7 family members in XRN2-depleted HUH7 tumors compared to control tumors.

**g.** Representative images of mice injected subcutaneously with either scramble shRNA or XRN2 shRNA transfected U87-MG cells. Imaged and investigated for tumor status after 45 days of injection.

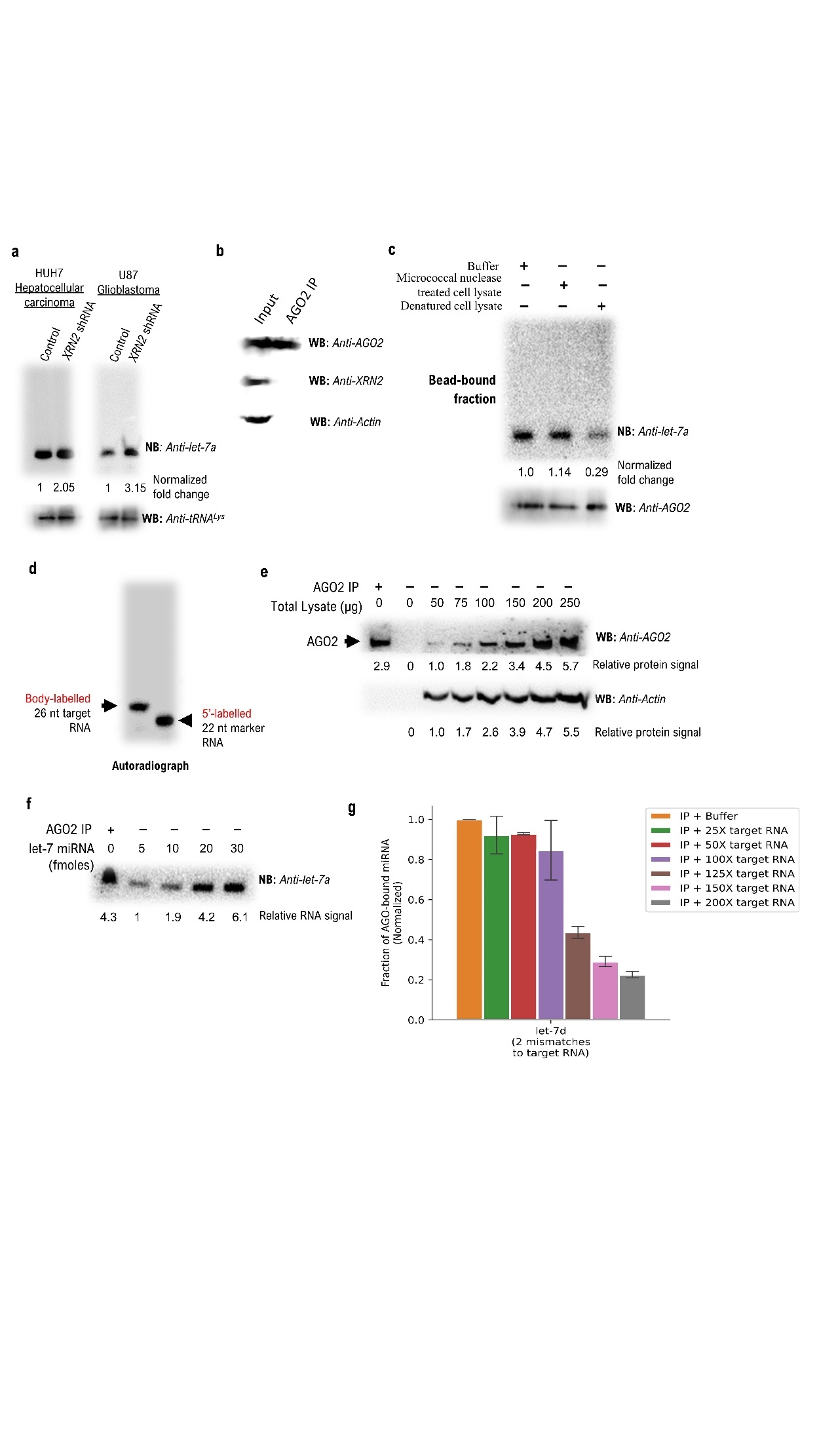

**Figure S7. Critical parameters of AGO2 IP and controls for miRNA release assays.**

**a.** Immunoprecipitation using anti-AGO2 antibody from total cell lysate of the indicated samples reveal increased signal of let-7a in AGO2 of XRN2-depleted HUH7 and U87 cells. One half of each immunoprecipitate was subjected to northern probing, the other half to anti-AGO2 western probing and served as loading control.

**b.** Co-immunoprecipitation experiment indicated that interaction between XRN2 and AGO2 is not detectable in A549 cells, at the limit of detection of the employed western probing.

**c.** miRNA release assay with immunoprecipitated AGO2 from A549 cells was performed with the indicated samples. Pre-treatment of the cell lysate with a denaturing agent (urea) result in a complete loss of the miRNA releasing capability of the lysate. Denaturing agent was removed from the cell lysate by a single-step dialysis prior to its use in the ‘miRNA release assay’.

**d.** Autoradiograph confirms size and intactness of target RNA used for release assays, resolved alongside a labelled marker RNA of known size (as indicated).

**e.** Anti-AGO2 western probing performed using different amounts of total cell lysate indicates that recovered AGO2 in the immunoprecipitated sample is equivalent to the amount present in ~75 µg of total cell lysate (densitometric quantification).

**f.** Northern detection of varying amounts of synthetic let-7a indicates that the amount of let-7a recovered from AGO2 IP is equivalent to ~20 femtomoles (fmol).

**g.** TaqMan analyses reveal that let-7d miRNA is released from immunoprecipitated AGO2 from A549 cells upon treatment with increasing concentrations of a target RNA (perfectly complementary to let-7a, but with 2 mismatches for let-7d), albeit with lower efficiency compared to an assay employing a perfectly complementary target for a given miRNA (compare with **Fig. 5d)**.

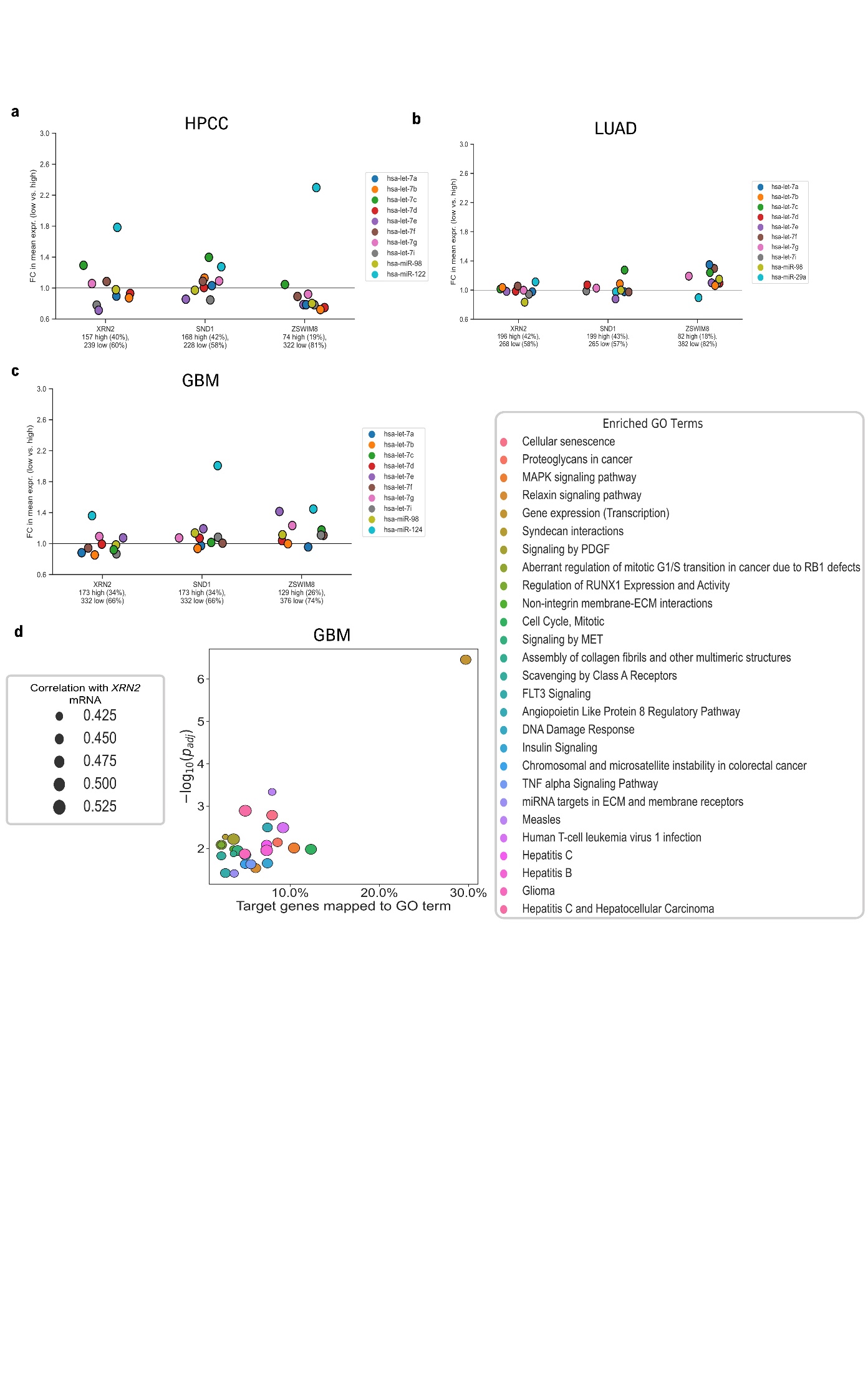

**Figure S8. Comparative analyses of the impacts of XRN2-mediated regulation in the TCGA-HPCC, TCGA-LUAD and TCGA-GBM complete datasets.**

**a.** Comparative analyses of mean fold changes of let-7 family members and liver-specific miR-122 in the complete TCGA-HPCC dataset with respect to low and high expression groups for *XRN2*, *SND1*, and *ZSWIM8* mRNAs. *SND1* and *XRN2* mRNA levels have comparable negative influence on levels of let-7 family members as well as miR-122.

**b.** Comparative analyses of mean fold changes of let-7 family members and miR-29a, in the complete TCGA-LUAD dataset with respect to low and high expression groups for *XRN2*, *SND1*, and *ZSWIM8* mRNAs. mRNA levels of *ZSWIM8* has the most prominent negative influence on levels of most let-7 family members, followed by *SND1* and *XRN2* mRNA levels. Only high *XRN2* mRNA levels appear to negatively influence levels of miR-29a in the complete dataset.

**c.** Comparative analyses of mean fold changes of let-7 family members and miR-124, in the complete TCGA-GBM dataset with respect to low and high expression groups for *XRN2*, *SND1*, and *ZSWIM8* mRNAs. mRNA levels of *SND1* and *ZSWIM8* have comparable negative influence on levels of let-7 family members, with XRN2 mRNA levels having a negative influence on let-7e and let-7g. High *XRN2* and *ZSWIM8* mRNA levels exert comparable negative influence on the levels of miR-124.

**d.** Gene Ontology (GO) Term Enrichment analysis indicate that the most enriched GO terms for the genes positively correlated with *XRN2* mRNA expression in the TCGA-GBM dataset, again include multiple signalling pathways, out of which “gene expression (transcription)”, “cellular senescence”, and “mitotic cell cycle” are some of the most enriched terms. Of note, the terms are largely shared with the GO Term Enrichment analysis plot for TCGA-HPCC, and there are cross-enrichment terms related to cancer and viral infection with Figure 6C, indicating that these affected pathways are preserved across different cell lineages.

**Supplementary Table 1**

Status of the validated let-7 targets derived from the mRNA-seq data presented in Fig. 2a, c, and Fig. S3a, b.

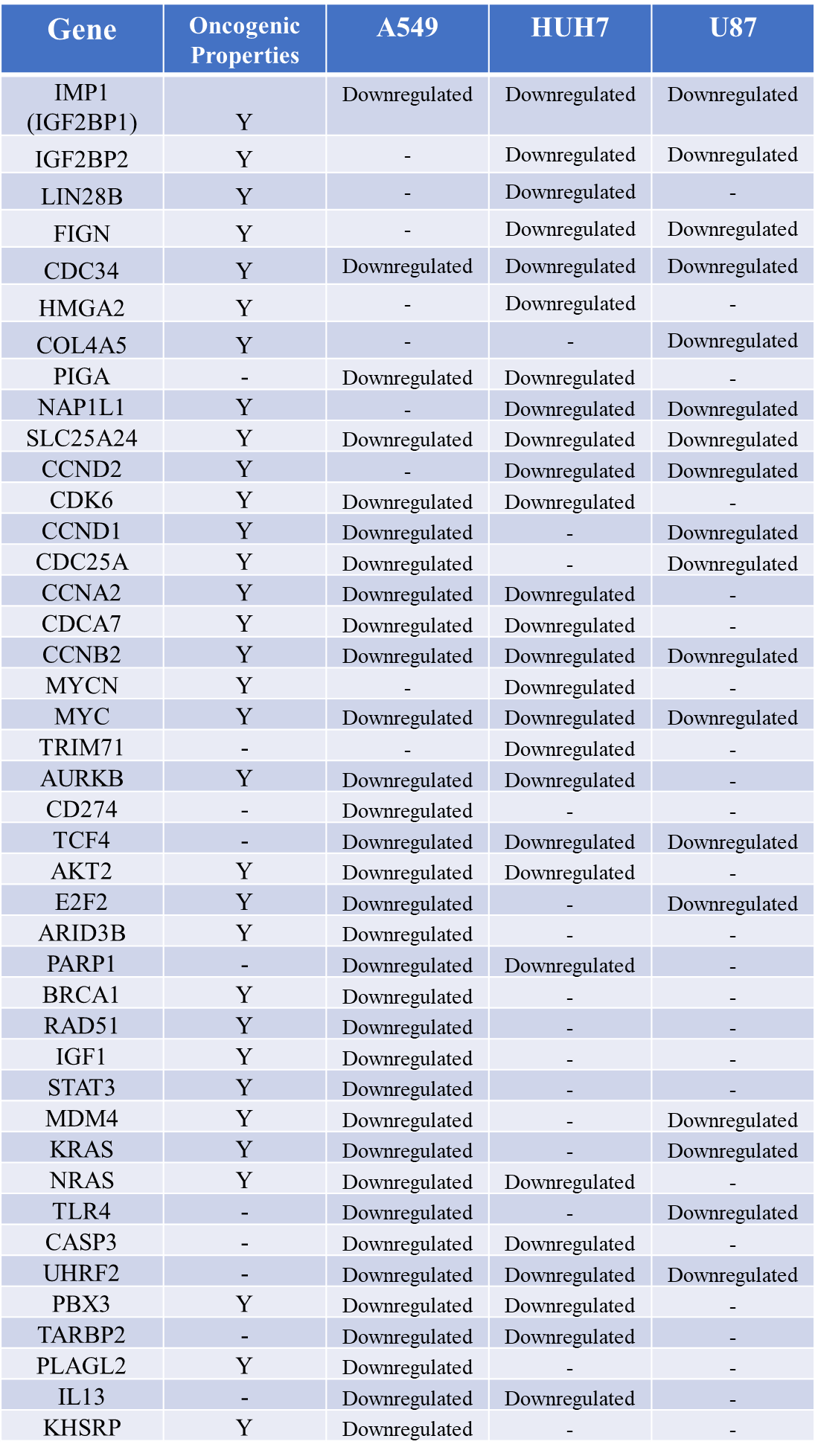

**Y – Yes.**

**Materials and Methods**

**Cell culture**

HEK293T, A549, MDA-MB-231, MCF-7, HUH7, HEPG2, LN229, U87-MG, and HeLa cells were cultured in Dulbecco's modified Eagle's medium (DMEM) supplemented with 10% fetal bovine serum (FBS) and 1% penicillin-streptomycin at 37 °C in 5% CO_2_.

**Experimental mice model**

Experiments were performed in Athymic Nude female mice (5–6-week-old) following the approval by the Institute Ethical Committee for Animal Experimentation. The mice were kept in a 12h light and dark cycle, fed *ad libitum* with a normal diet and the experiments were done during the light phase of the cycle.

**Lentivirus mediated knockdown**

Virus was produced in HEK293T cells, and at 72 hr post transfection, the medium containing the shRNA lentiviruses was collected and filtered using 45 μm filters. Transduction in desired cell line was carried out in the presence of 10 μg/mL of polybrene (Sigma; CAS Number: 28728-55-4) followed by selection with 2 μg/mL puromycin (Sigma; CAS Number: 58-58-2). Post selection, cells were harvested at 60, 72, and 96 hr timepoints and tested for knockdown efficiency.

TRC shRNA plasmids against *XRN2* were purchased from TRC genome-wide shRNA library (Sigma-Aldrich). Empty pLKO.1-Puro plasmid was used as a control in all shRNA knockdown experiments. Lentiviral packaging vectors pMD2.G (VSV-G envelope, 12259) and psPAX2 (packaging, 12260) were used for virus generation.

**Transfection of let-7a antagomirs**

MISSION Synthetic miRNA Inhibitors from Sigma-Aldrich were used to inhibit let-7a function. 10 nm of let-7a inhibitor (Sigma, HSTUD0001) and control inhibitor (Sigma, NCSTUD001) were transfected into the cells seeded in 6-well plates. Transfection was performed using Lipofectamine™ 3000 Transfection Reagent (Invitrogen™) (Cat No.: L3000001) according to the manufacturer's protocols.

**Clonogenic/ Colony formation assay**

Cells were plated (1000 cells/well) and grown for 2 weeks replacing the media every 3 days. Colonies were fixed in chilled methanol followed by staining with crystal violet (0.5%) for 20 minutes. Colonies were quantified post imaging of the 6-well plate using ImageJ (NIH)^9^.

**Wound healing/ Scratch assay**

Migration was performed using scratch assay. Briefly, 0.25 x 10^6^ cells/well were plated in a 12-well plate. After 24 hr, scratch was made and DMEM media was added without FBS. Images were taken at 0 and 24 hr post scratch creation. The distance across a scratch was measured using ImageJ (NIH)^9^, and the gap measured at the 0^th^ hr was set to 100%.

**Subcutaneous injections in mice**

100 μl of cell suspension (5*10^6^ cells) was injected subcutaneously into right flank of nude mice (n = 5). The tumor diameters were measured every 3 days using Vernier calipers. Tumor volumes were determined using the formula: V = D*d^2^/2 (where “V” refers to tumor volume, “D” is largest diameter and “d” is smallest diameter).

**Intra-cranial injection in mice**

Cells were harvested in incomplete DMEM, 4*10^5^ cells (per mouse) were injected intracranially (in the right cerebrum, 3 mm deep) in each animal using a stereotaxic apparatus. The animals were imaged on the 3rd day after injection and subsequently, every 5 to 6 days until the end of the experiment.

***In vivo* imaging**

*In vivo* imaging was done for bioluminescence with the Perkin Elmer IVIS Spectrum by using mild gas anesthesia (using isoflurane) for the animals.

**Hematoxylin and Eosin staining**

Brains from perfused mice were paraffin embedded and sectioned using a microtome (5μ sections). Sections were mounted on glass slides and removed from paraffin, rehydrated and stained with Harris Hematoxylin for nuclear staining and Eosin Y solution for cytoplasmic staining. The sections were then mounted using DPX mounting medium and imaged at 0.8X using a Lawrence and Mayo digital microscope.

**RNA isolation, Northern blotting, RT-qPCR**

Total RNA from cells was extracted using TRIzol reagent (Ambion) as per the manufacturer's instructions. Northern detection of endogenous miRNA (30 µg/sample) was performed as described in (1). 5’- radiolabelled DNA oligos were used as probes. The hybridization for let-7a miRNA was carried out at an elevated temperature of 40°C to minimize the binding of the probe to unintended let-7 sisters.

For RT-qPCR, 2 µg of total RNA of the respective samples were reverse transcribed using SuperScript™ III Reverse Transcriptase (Invitrogen) in 20 µl reactions containing 5 mM DTT, 0.5 mM dNTPs, 2.5 µM oligo(dT)_20_ and 1 µl of enzyme, in accordance with the manufacturer’s instructions. 1.5 µl of each of the reverse-transcription reactions were amplified with PowerUp^TM^ SYBR Green Master Mix (Applied Biosystems) in a total volume of 10 µl using specific forward and reverse primers at a concentration of 0.5 µM each. All reactions were carried out in an Applied Biosystems QuantStudio™ 5 Real-Time PCR System and analysed using ΔΔCt method. The primers used are furnished in the supplementary table. The experimental values, representing means (±SEM) from three independent biological replicates, were compared to the expression levels of the corresponding control samples. The ratio of pre-miRNA transcripts to pri-miRNA transcripts were determined as described in (13).

For quantification of miRNAs, TaqMan reverse transcriptase kits were used, followed by a TaqMan universal PCR mix per the manufacturer's instructions (Applied Biosystems). All reactions were carried out in Applied Biosystems QuantStudio™ 5 Real-Time PCR System and analysed using ΔΔCt method. Cycles were set as per the manufacturer's instructions.

**Luciferase reporter assay**

Cells were seeded in 6-well plates and co-transfected with the reporter Renilla luciferase construct and Firefly luciferase construct (as control), using lipofectamine according to the manufacturer’s instructions. 48 h after transfection, cells were harvested, and luciferase activity in the cells was analysed using a Dual-Luciferase Reporter Assay System (Promega).

**Argonaute (AGO2) immunoprecipitation and Western blotting**

Endogenous Argonaute protein AGO2/ AGO2 complexes were immunoprecipitated using anti-EIF2C2 (anti-AGO2) mouse IgG; monoclonal antibody, Catalog #: H00027161-M01; Abnova) and Protein G-Sepharose 4 FastFlow (GE Healthcare) as described in (2).

For Western blotting, after SDS-PAGE, resolved proteins were transferred to polyvinylidene difluoride (PVDF) membrane, followed by blocking and probing with the following primary antibodies at 4°C for 12 hr – rabbit Anti-Argonaute-2 antibody (Abcam, 1:1000); rabbit anti-human XRN2 antibody (A301-103A, Bethyl Laboratories 1:1000); rabbit anti-XRN1 antibody (A300-443A, Bethyl Laboratories 1:2000). Detection was performed with Amersham^TM^ ECL^TM^ Prime Western Blotting Detection Reagent, and ImageQuant LAS 4000 Chemiluminescence Imager (GE Healthcare). Densitometric quantification of all Western Blots was performed using ImageJ (NIH)^9^.

**miRNA release assay**

The bead-bound anti-AGO2 immunoprecipitates (derivative of 1.5 mg of total lysate per reaction) were incubated with 1X assay buffer (AB; 20mM HEPES pH 7.56, 3 mM DTT, 7.5 mM MgCl_2_, 100 mM KCl), or 1$\times$ AB plus KCl (to a final concentration of about 1.0 M), or 75 µg of micrococcal nuclease/EGTA-treated control or XRN2-depleted cell lysates, at 37°C for 30 minutes. After further recovery, the beads were split into two halves. RNA was extracted from one half for performing Northern blotting or TaqMan analysis; the other half was boiled in SDS sample buffer and subjected to SDS–PAGE and western blotting with Anti-AGO2 antibody, to serve as a loading control and confirm the integrity of the proteins. RNA was also extracted from each of the above supernatant fractions and subjected to northern analysis to detect miRNAs released in the supernatant.

The cell lysates employed in the miRNA release assays were pre-treated with micrococcal nuclease (MN, NEB, 2.0 µl (4000 Gels Units)/100 µg of lysate) for 10 minutes at 37°C, followed by addition of EGTA to a final concentration of 7.5 mM. MN pre-treatment was done to digest all endogenous RNAs from the lysate, thus ruling out either any role played by the endogenous RNA in ‘miRNA release’ or possibility of detection of endogenous RNA in subsequent northern probing. Excess EGTA was used to terminate the MN treatment through chelation of Ca^2+^. Further, MN-treated cell lysates were appropriately diluted with 1X assay buffer and used for subsequent assays. Following incubation of the bead-bound immunoprecipitated samples with the lysates, supernatant fractions were removed, and re-recovered bead-bound fractions were phenol-chloroform extracted and alcohol precipitated in the presence of glycogen (Roche, 20 µg/ reaction). The recovered RNAs were suspended in Nuclease-free water (Ambion) for TaqMan analysis as stated below. For northern analysis, the recovered RNAs were suspended in formamide gel loading buffer, heated at 65°C for 5 minutes, centrifuged briefly, and subjected to urea-PAGE analysis followed by northern probing as per the conditions stated above.

For target RNA-mediated release assay, the bead-bound immunoprecipitates were incubated with 1X AB having different concentrations of the *in vitro* transcribed target RNA at 37°C for 30 minutes. Post incubation, beads were re-recovered (after appropriate washes with 1$\times$ AB). The re-recovered beads were split into two halves. RNA was extracted from one half and re-suspended in Nuclease-free water (Ambion) for TaqMan analysis as stated above; the other half was subjected to SDS–PAGE and Western analysis.

For denaturation, the total cell lysate was subjected to denaturation buffer (50 mM Tris, 50 mM NaCl, 10 mM β-mercaptoethanol, 8 M Urea, pH 8.0), then left on a rotator for 20 minutes at RT, and centrifuged (30,000 g, 4°C, 45 minutes). The supernatant was collected, and denaturants were dialysed out using large volume of dialysis buffer in a single-step (Dialysis buffer: 50 mM Tris, 1 mM EDTA, 3.0 mM MgCl_2_, 50 mM NaCl, pH 8.0). Total protein concentration was measured after dialysis by using the Bradford method and used accordingly for the assay.

**Nucleocytoplasmic fractionation of cells**

Sub confluent cells grown on 10-cm plates, were subjected to mild lysis in hypotonic buffer (20 mM Tris⋅Cl, pH 7.5, 10 mM KCl, 0.5% NP-40 (Nonidet P-40), 3 mM DTT, 3 mM MgCl2, 0.2 mM EDTA, 1X PIC) for 15 min at 4°C. The cytoplasmic fraction was obtained as supernatant after centrifugation at 2,000 × g for 5 min. The pellet was washed twice in hypotonic buffer and extracted with hypertonic buffer (20 mM Tris⋅Cl, pH 7.5, 600 mM KCl, 3 mM DTT, 0.2 mM EDTA, 1X PIC), yielding the nuclear fraction. Both cytoplasmic and nuclear fractions were analyzed by Western blotting.

**Nucleocytoplasmic Fractionation of total RNA**

Cells growing in 10 cm dishes were rinsed twice with ice-cold 1X PBS, harvested in 1 mL ice-cold 1X PBS by scraping, and centrifuged at 1,000 rpm for 10 minutes. Cell pellets were

resuspended by gentle pipetting in 200 µL lysis buffer A (10 mM Tris (pH 8.0), 140 mM NaCl,

1.5 mM MgCl2, 0.5% NP-40), incubated on ice for 5 minutes with intermittent taping, and then centrifuged at 1,000 X g for 3 minutes at 4°C. The supernatant, containing the cytoplasmic fraction, was collected, and subjected immediately to 1 mL Trizol for RNA purification. Nuclear pellets underwent two additional washes with lysis buffer A and a final wash with lysis buffer B (lysis buffer A containing 1% Tween-40 and 0.5% deoxycholic acid). Purified nuclear pellets were then resuspended in 1 mL Trizol. For northern blotting, 1/3 of the total yield of RNA from nuclear and cytoplasmic fractions was loaded to allow comparison of equal cell equivalents.

***In vitro* transcription of target RNA**

Perfectly complementary let-7a target RNA was transcribed from DNA cassettes using a MEGAshortscript^TM^ T7 transcription kit (Ambion) in presence of α-^32^P-UTP, as per the supplier’s instructions. The DNA cassettes were prepared by annealing of appropriate forward and reverse primers (refer oligo sequence section) followed by fill-in reactions using Klenow DNA Polymerase (NEB). Double stranded DNAs of appropriate length were gel purified and directly used as template for *in vitro* transcription. The gel purified products were also PCR amplified, cloned and sequence confirmed. After transcription reaction was over, the resulting mature let-7a target transcript was size purified by resolving the reaction products on 7 M urea/10% PAGE.

**α-Amanitin treatment**

Cells were seeded in each well of a 6-well plate and incubated for 16 hours to reach 90% confluency. Post incubation, cells were treated with α-Amanitin (10 µg/ml; Sigma, A2263) and harvested at the time points indicated in the results.

**mRNA cDNA library cloning**

DNase-treated total RNA with RIN > 8 was used to prepare strand-specific RNA libraries. Firstly, poly(A)+ RNA was isolated from total RNA using NEBNext® Poly(A) mRNA Magnetic Isolation Module (NEB). Then strand-specific RNA-seq libraries were prepared using the NEBNext® Ultra™ II Directional RNA Library Prep Kit (NEB) according to the manufacturer’s instructions.

**mRNA sequencing analysis**

Reads were adapter-trimmed using fastp v0.20.0^16^ (fastp --in1 "$sample_R1.fq.gz" --in2 "$sample_R2.fq.gz" --out1 "$sample_R1.trim.fq.gz" --out2 "$sample_R2.trim.fq.gz" -q 20 -u 10 --detect_adapter_for_pe –correction). Trimmed reads were then aligned to the human genome (GRC hg38.p13 from the Ensembl database version 106^17^) using the HISAT2^18^ splice-aware aligner with default settings. The samtools^19^ collate-fixmate-markdup pipeline was used to eliminate optical duplicates in the resulting .bam file, and reads were assigned to features in the canonical geneset using Subread featureCounts 2.0.3^20^ (featureCounts -a GRCh38.refseq_annotation.gtf.gz -p --countReadPairs -C -M -O --fracOverlap 0.01 --fraction --ignoreDup --largestOverlap -o "$output" "${aligned_files[@]}"). The fractional read counts output from featureCounts was then analyzed using DESeq2^21^ to identify differentially expressed genes (DEGs), defined as genes with $FC\leq0.7 \text{or} FC\geq1.3$, and $p_{adj}\leq0.1$. Genes targeted by let-7 family miRNAs were determined by predictions from TargetScan Human release 7.2^8^. Expression levels of let-7 target and other mRNAs of interest were normalized to Transcripts per Million (TPM) across all samples, and each mRNA was individually scaled as maximum expression to unity across samples for purposes of heatmap generation. DESeq2 analysis output was directly used to generate MA/ Bland-Altman or volcano plots.

**Bioinformatic analysis**

miRNA isoform and mRNA expression quantification files were downloaded for every solid tumour sample under the following datasets: TCGA-HPCC (hepatocellular carcinoma), TCGA-GBM (glioblastoma) and TCGA-LUAD (lung adenocarcinoma). Samples having any deleterious mutations associated with the *XRN2* gene (from the SIFT^3^ or PolyPhen^4^ single-nucleotide polymorphism (SNP) genotype-phenotype association databases) were discarded. For each dataset, samples were divided into high and low *XRN2* expression categories depending on if their *XRN2* mRNA levels were above or below the median for the dataset, respectively. Differences in expression of let-7 target mRNAs, heatmaps and volcano plots of let-7 family miRNA expression were quantified and visualized using a custom Python script employing Python 3.8.2 and Pandas 1.2.4 with Numpy 1.20, SciPy 1.6.3, Matplotlib 3.4.1 and Seaborn 0.11.1. Unless otherwise specified, all p-values were calculated using the Student’s t-test with the Bonferroni correction for multiple testing (p-value cut-offs unless otherwise specified: $* \Rightarrow0.1\leq p<0.01$, $** \Rightarrow0.001<p\leq0.01$, $*** \Rightarrow0.0001<p\leq0.001$, $**** \Rightarrow p\leq0.0001$, $ns\Rightarrow p>0.1$). Statistical functions used were from the scipy.stats package with default settings. All expression values were normalized to Transcripts Per Million (TPM) from FPKM (mRNA) or RPM (miRNA) before fold change calculations were performed or any plots were generated.

**CDF analysis**

CDF analysis of the mRNA-seq data was performed as described in reference^15^.

**‘Representative Subset’ determination:**

Random uniform sampling of 100 samples from the relevant TCGA dataset was performed repeatedly while minimizing the combined mean fold change of let-7c and let-7g for high vs. low *XRN2* mRNA expression through minimization of the Euclidean distance of the fold change vector from 0. This serves to maximize the negative (inverse) fold change, as the maximum possible negative (inverse) fold change would be 0. In each run, the best amongst 20 trials was recorded. Amongst the best of 10 such runs, the sample subset displaying the greatest combined inverse correlation of let-7c and let-7g with *XRN2* mRNA levels was chosen as the ‘representative subset’ for further comparative analyses, including survival analysis and the determination of miRNAs possessing an inverse relationship with *XRN2* mRNA levels (Submitter IDs of TCGA ‘representative subset’ samples have been furnished below).

**Comparison of negative regulation of let-7 family members by *XRN2*, *SND1* and *ZSWIM8*:**

The high and low expression groups for *XRN2*, *SND1*, and *ZSWIM8* mRNA were established in the ‘representative subset’ and complete dataset for TCGA-HPCC, TCGA-GBM, and TCGA-LUAD, on the basis of median expression as described before, and the fold change in the mean expression of each let-7 family member between the low and high expression groups for *XRN2*, *SND1*, and *ZSWIM8* mRNAs were calculated.

**Survival Analysis:**

Kaplan-Meier survival analysis for the samples in each category was performed on the ‘representative subset’ from the respective TCGA dataset using the survival analysis endpoint of the TCGA web service (<https://api.gdc.cancer.gov/analysis/survival>), and significance of the difference in survival was established using a Chi-Square test with 1 degree of freedom (2 categories).

**Gene Ontology Term Enrichment Analysis:**

Gene Ontology term enrichment analysis was performed using the g:OSt tool from the g:Profiler web service (<https://biit.cs.ut.ee/gprofiler/gost>)^5^, version e104_eg51_p15_3922dba, using the KEGG^6^ and Reactome+7 pathway databases with all other settings at the recommended defaults. let-7 target genes were identified using TargetScan Human release 7.2^8^. The term intersections were downloaded as a CSV file from g:OSt and used in combination with the TCGA data to generate the plot in Figure 6B using the custom Python script.

All code and scripts for this project are available at the GitHub link: <https://github.com/tamchow/xrn2_let7_human_cancer>.

**RT-qPCR Primers**

| **Name** | **Primers** | |
| --- | --- | --- |
| VIM | For - 5‘-CCTGCAGGAGGCAGAAGAATG-3’  Rev - 5’-GTTCCAGGGACTCATTGGTTCC-3’ | |
| CDH1 | For - 5’-GTGCCTGAGAACGAGGCTAA-3’  Rev - 5’-CTGCATCTTGCCAGGTCCTT-3’ | |
| KRAS | For - 5’-TCGACACAGCAGGTCAAGAGGA-3’  Rev - 5’-ATCCTCCACTCTCTGTCTTGTCTT-3’ | |
| HMGA2 | For - 5’-GGAAATGGGACAATCTACTACCAA-3’  Rev - 5’-CGGAGAAAGCCACACATAAGG-3’ | |
| CDC25A | For - 5′-CTACCTCCCACACTCCCAAG-3′  Rev - 5′-ACTTCTCTACTCCCCTCCGT-3′ | |
| PLK1 | For - 5’-GCACAGCACAGTGTCAATGCCTCCAAG-3’  Rev - 5’-GCCGTACTTGTCCGAATAGTCC-3’ | |
| KLK6 | For - 5′-GAAGCATAACCTTCGGCAAA-3′  Rev - 5′-GGGAAATCACCATCTGCTGT-3′ | |
| NRAS | For - 5’-AGACCAGACAGGGTGTTGAAG-3’  Rev - 5’-AAGTCAGGACCAGGGTGTCA-3’ | |
| SNAI1 | | For - 5’-CAGAGTTTACCTTCCAGCAGCC-3’  Rev - 5’-CTCATCTGACAGGGAGGTCAGC-3’ |
| CCND1 | | For - 5’-ATGCCAACCTCCTCAACGAC-3’  Rev - 5’-CGTGGGTCTGGGCAACAAGTT-3’ |
| ACTB | | For – 5’-CTGGAACGGTGAAGGTGACA-3’  Rev – 5’-AAGGGACTTCCTGTAACAATG-3’ |
| ATP5G1 | | For – 5’-CCAGACGGGAGTTCCAGAC-3’  Rev – 5’-GACGGGTTCCTGGCATAGC-3’ |
| pri-let-7a-1 | | For – 5’-GATTCCTTTTCACCATTCACCCTGGATGTT-3’  Rev – 5’- TTTCTATCAGACCGCCTGGATGCAGACTTT-3’ |
| pri-let-7g | | For – 5’-GTTCCTCCAGCGCTCCGTT-3’  Rev – 5’-CCATTACCTGGTTTCCCAGAGA-3’ |
| pri-let-7i | | For – 5’-GTGCCTCCCCGACACCAT-3’  Rev – 5’-GTGAAACTAACGGTTTCCGTGGT-3’ |
| pre-let-7a-1 | | For – 5’-TGAGGTAGTAGGTTGTATAGTT-3’  Rev – 5’-TCCCAGTGGTGGGTGTGACCCTAAA-3’ |
| pre-let-7g | | For – 5’-GGCAAGGCAGTGGCCTGTACAGTT-3’  Rev – 5’-TGAGGTAGTAGTTTGTACAGTT-3’ |
| pre-let-7i | | For – 5’-TGAGGTAGTAGTTTGTGCTGTT-3’  Rev – 5’-AGCAAGGCAGTAGCTTGCGCAG-3’ |

**Northern primers**

mature let-7a: 5’ – AACTATACAACCTACTACCTCA – 3’

tRNA^lys^: 5’ – CTGATGCTCTACCGACTGAGCTATCCGGGC – 3’

mature miR-21: 5’ – TCAACATCAGTCTGATAAGCTA – 3’

***In vitro* transcription primers**

Perfectly complementary let-7a target RNA

For – 5’ – GTAATACGACTCACTATAGGG – 3’

Rev – 5’ – CCTGAGGTAGTAGGTTGTATAGTTCCCTATAGTGAG – 3’

**Submitter IDs of TCGA ‘representative subset’ samples**

| **TCGA-HPCC** | **TCGA-GBM** | **TCGA-LUAD** |
| --- | --- | --- |
| TCGA-DD-A39Z | TCGA-VV-A829 | TCGA-75-6206 |
| TCGA-UB-A7MA | TCGA-DU-A5TT | TCGA-J2-8192 |
| TCGA-MI-A75I | TCGA-HT-7474 | TCGA-86-8074 |
| TCGA-G3-A5SK | TCGA-E1-A7YI | TCGA-05-4415 |
| TCGA-WD-A7RX | TCGA-S9-A6WL | TCGA-44-7660 |
| TCGA-G3-A7M5 | TCGA-HT-7695 | TCGA-55-A491 |
| TCGA-DD-AAVV | TCGA-DU-5855 | TCGA-05-4420 |
| TCGA-DD-AAVP | TCGA-VM-A8CB | TCGA-69-A59K |
| TCGA-ZP-A9D4 | TCGA-CS-5393 | TCGA-71-8520 |
| TCGA-CC-5258 | TCGA-FG-A87N | TCGA-44-3919 |
| TCGA-W5-AA2G | TCGA-R8-A6YH | TCGA-78-7540 |
| TCGA-DD-AACW | TCGA-DB-A4XH | TCGA-55-7574 |
| TCGA-KR-A7K0 | TCGA-TQ-A7RF | TCGA-55-7227 |
| TCGA-UB-A7MF | TCGA-S9-A7IY | TCGA-05-4397 |
| TCGA-CC-5261 | TCGA-DB-A75K | TCGA-MP-A4SW |
| TCGA-DD-AAD2 | TCGA-DU-6394 | TCGA-L4-A4E5 |
| TCGA-DD-A4NO | TCGA-HT-7881 | TCGA-55-7815 |
| TCGA-CC-A7II | TCGA-QH-A6X4 | TCGA-05-4432 |
| TCGA-DD-A4NR | TCGA-E1-A7YU | TCGA-44-A479 |
| TCGA-DD-AAW2 | TCGA-S9-A89Z | TCGA-78-7153 |
| TCGA-BC-A69I | TCGA-WY-A85C | TCGA-MP-A5C7 |
| TCGA-UB-A7MC | TCGA-TM-A7C5 | TCGA-05-4422 |
| TCGA-DD-A114 | TCGA-R8-A73M | TCGA-64-1678 |
| TCGA-W5-AA2T | TCGA-E1-5302 | TCGA-75-7025 |
| TCGA-DD-AADF | TCGA-DB-A64X | TCGA-69-7765 |
| TCGA-ED-A7PX | TCGA-DU-6393 | TCGA-05-4426 |
| TCGA-ZP-A9D0 | TCGA-QH-A6X8 | TCGA-75-7031 |
| TCGA-DD-AAE9 | TCGA-DU-A5TY | TCGA-50-5044 |
| TCGA-5C-AAPD | TCGA-TQ-A7RS | TCGA-83-5908 |
| TCGA-MR-A8JO | TCGA-FG-8186 | TCGA-97-8175 |
| TCGA-DD-AAW3 | TCGA-DU-6396 | TCGA-J2-A4AG |
| TCGA-G3-A5SJ | TCGA-DU-5847 | TCGA-44-2661 |
| TCGA-G3-AAV3 | TCGA-IK-8125 | TCGA-49-6745 |
| TCGA-XR-A8TD | TCGA-P5-A737 | TCGA-44-A47B |
| TCGA-DD-A11C | TCGA-HT-8019 | TCGA-05-4418 |
| TCGA-CC-A8HU | TCGA-VW-A8FI | TCGA-05-4427 |
| TCGA-2Y-A9H9 | TCGA-FG-8182 | TCGA-50-5941 |
| TCGA-DD-A4NB | TCGA-DH-5142 | TCGA-75-6207 |
| TCGA-DD-A1EJ | TCGA-HW-7490 | TCGA-67-3770 |
| TCGA-CC-A7IF | TCGA-QH-A6CU | TCGA-75-6203 |
| TCGA-G3-A3CK | TCGA-HT-8015 | TCGA-55-8510 |
| TCGA-2Y-A9GS | TCGA-VM-A8CA | TCGA-55-6979 |
| TCGA-W5-AA2U | TCGA-DU-A7T6 | TCGA-55-A57B |
| TCGA-ZU-A8S4 | TCGA-HT-8558 | TCGA-05-4430 |
| TCGA-G3-AAV1 | TCGA-TM-A84F | TCGA-55-6712 |
| TCGA-BD-A2L6 | TCGA-S9-A7R1 | TCGA-97-A4M0 |
| TCGA-NI-A4U2 | TCGA-E1-5322 | TCGA-69-8254 |
| TCGA-DD-A115 | TCGA-P5-A731 | TCGA-55-6984 |
| TCGA-CC-5262 | TCGA-FG-6688 | TCGA-44-A4SS |
| TCGA-DD-A4NV | TCGA-VM-A8CD | TCGA-55-1596 |
| TCGA-G3-A3CH | TCGA-HT-7604 | TCGA-97-A4M5 |
| TCGA-WJ-A86L | TCGA-TQ-A7RV | TCGA-44-7667 |
| TCGA-DD-AAEK | TCGA-HT-A5RA | TCGA-55-8621 |
| TCGA-BC-A10R | TCGA-S9-A6WG | TCGA-91-A4BD |
| TCGA-FV-A2QR | TCGA-S9-A7R3 | TCGA-55-6543 |
| TCGA-DD-AACU | TCGA-HT-A61B | TCGA-95-7043 |
| TCGA-DD-AACN | TCGA-FG-A4MY | TCGA-49-AAQV |
| TCGA-DD-AACY | TCGA-HT-7873 | TCGA-97-A4M2 |
| TCGA-ZH-A8Y4 | TCGA-DU-A7TD | TCGA-86-8358 |
| TCGA-W5-AA39 | TCGA-DB-5279 | TCGA-L9-A50W |
| TCGA-DD-AAD3 | TCGA-EZ-7264 | TCGA-86-A4JF |
| TCGA-DD-A1EL | TCGA-HT-7857 | TCGA-MN-A4N5 |
| TCGA-4G-AAZO | TCGA-CS-6666 | TCGA-71-6725 |
| TCGA-FV-A2QQ | TCGA-QH-A6CV | TCGA-78-7160 |
| TCGA-G3-A7M9 | TCGA-S9-A7IS | TCGA-55-8091 |
| TCGA-DD-AAVU | TCGA-TM-A84L | TCGA-78-7539 |
| TCGA-2Y-A9GX | TCGA-DU-A7TI | TCGA-55-8208 |
| TCGA-XR-A8TC | TCGA-DB-A4XB | TCGA-55-A493 |
| TCGA-UB-A7MD | TCGA-DU-A6S7 | TCGA-38-4631 |
| TCGA-DD-AADM | TCGA-QH-A65Z | TCGA-91-6840 |
| TCGA-O8-A75V | TCGA-HT-8105 | TCGA-97-7547 |
| TCGA-DD-AAEI | TCGA-FG-8189 | TCGA-NJ-A7XG |
| TCGA-W5-AA33 | TCGA-HT-A74K | TCGA-99-7458 |
| TCGA-RC-A6M3 | TCGA-DU-7300 | TCGA-93-8067 |
| TCGA-2Y-A9H1 | TCGA-E1-5304 | TCGA-64-5815 |
| TCGA-UB-AA0V | TCGA-HT-7609 | TCGA-05-4417 |
| TCGA-3X-AAVB | TCGA-TQ-A7RG | TCGA-86-7954 |
| TCGA-DD-AAE3 | TCGA-E1-A7YS | TCGA-55-6987 |
| TCGA-G3-AAV6 | TCGA-DU-7302 | TCGA-86-8359 |
| TCGA-W5-AA2Z | TCGA-DB-A64Q | TCGA-L9-A443 |
| TCGA-FV-A3I0 | TCGA-CS-4943 | TCGA-86-7955 |
| TCGA-ES-A2HS | TCGA-HT-7478 | TCGA-55-8301 |
| TCGA-BC-A10U | TCGA-FG-7641 | TCGA-44-6776 |
| TCGA-BC-4073 | TCGA-HW-8319 | TCGA-05-4433 |
| TCGA-ED-A66X | TCGA-QH-A65S | TCGA-50-6595 |
| TCGA-DD-A73E | TCGA-HT-7479 | TCGA-67-6215 |
| TCGA-CC-A3M9 | TCGA-HT-7856 | TCGA-64-5781 |
| TCGA-ZS-A9CE | TCGA-HT-8564 | TCGA-78-8662 |
| TCGA-DD-A1EH | TCGA-DH-A7UV | TCGA-55-7281 |
| TCGA-DD-AA3A | TCGA-E1-A7Z6 | TCGA-55-8615 |
| TCGA-CC-A1HT | TCGA-E1-A7YM | TCGA-55-7907 |
| TCGA-W5-AA34 | TCGA-FG-7638 | TCGA-49-4486 |
| TCGA-CC-5263 | TCGA-DU-A5TW | TCGA-55-6968 |
| TCGA-DD-AACH | TCGA-DH-A669 | TCGA-L9-A7SV |
| TCGA-G3-A25U | TCGA-TM-A7C4 | TCGA-55-7725 |
| TCGA-MI-A75E | TCGA-DU-6402 | TCGA-55-8302 |
| TCGA-FV-A495 | TCGA-DH-5144 | TCGA-44-8119 |
| TCGA-G3-A25V | TCGA-P5-A72W | TCGA-55-7995 |
| TCGA-DD-A1EI | TCGA-RY-A843 | TCGA-78-8655 |
| TCGA-3X-AAVE | TCGA-S9-A7J0 | TCGA-73-4659 |
